# PEtab.jl: Advancing the Efficiency and Utility of Dynamic Modelling

**DOI:** 10.1101/2025.04.30.651378

**Authors:** Sebastian Persson, Fabian Fröhlich, Stephan Grein, Torkel Loman, Damiano Ognissanti, Viktor Hasselgren, Jan Hasenauer, Marija Cvijovic

## Abstract

Dynamic models are useful to study processes ranging from cell signalling to cell differentiation. Common modelling workflows such as model exploration and parameter estimation are computationally demanding. The Julia programming language is a promising tool to address these computational challenges. To evaluate it we developed `SBMLImporter.jl` and `PEtab.jl,` a package for model fitting. `SBMLImporter.jl` was used to evaluate different stochastic simulators against `PySB` and `RoadRunner,` overall Julia simulators proved fastest. For Ordinary Differential Equations (ODE) models solvers, gradient methods, and parameter estimation performance were evaluated using PEtab benchmark problems. For the latter two tasks `PEtab.jl` was compared against `pyPESTO,` which employs the high-performance AMICI library. Guidelines for choosing ODE solver were produced by evaluating 31 ODE solvers for 29 models. Further, by leveraging automatic differentiation `PEtab.jl` proved efficient and, for up to medium-sized models, was often at least twice faster than `pyPESTO`, showcasing how Julia’s ecosystem can accelerate modelling workflows.

## Introduction

Understanding the complex nature of dynamic processes such as nutrient signalling, cell division, and apoptosis is one of the central aims in systems biology and pharmacology [25, 26]. A powerful tool to help achieve this goal is dynamic modelling, where chemical reactions are modelled via stochastic or deterministic rate equations [28]. To date, dynamic models have been used to study a range of processes, from small receptor networks [2] over stochastic gene expression [62] to large cancer signalling pathways [8].

Models with stochastic rate equations correspond to jump problems where reactions trigger changes (jumps) in species amounts, whereas, for deterministic rates, the model can be formulated as a system of Ordinary Differential Equations (ODEs) [15]. In either case, most models contain unknown parameters, such as reaction rate constants. Thereby, to understand a model’s characteristics it must be simulated for many parameter sets to validate if it can capture experimental observations. Since most models lack closed-form solutions, numerical simulation methods are needed. Hence, efficient modelling requires fast and flexible software.

For simulating and training stochastic and deterministic models, established MATLAB and Python packages exist. Tools like `RoadRunner` [60] and `PySB` [33] facilitate simulations, while parameter estimation for stochastic models can be performed with methods such as Approximate Bayesian Computation [27]. For deterministic ODE models, mature packages like `Data2Dynamics` and `pyPESTO,` allow for model training, identifiability analysis, and sensitivity analysis, with support for efficient gradient computations. However, for stochastic models, tasks like model training and identifiability are challenging, and despite being actively researched [4], require thousands to millions of model simulations. Furthermore, most packages for both stochastic and deterministic models suffer from the “two-language problem”, where performance-critical code is written in a high-performance language like C++, and the interface in a scripting language like Python, complicating package development and maintenance. This is also a key reason why many ODE packages lack support for automatic differentiation [1]. As a result, derivative computations partially rely on analytically derived formulas. This makes it challenging to perform non-standard tasks such as incorporating qualitative biological data [37, 51] or time-to-event data [6] into parameter estimation and performing automatic model structure discovery via scientific machine learning approaches [42].

The Julia programming language was recently proposed as a tool to meet the needs in system biology [3, 47]. For both stochastic and deterministic models, Julia’s ecosystem provides support for symbolic model pre-processing and state-of-the-art solvers [43, 32]. Additionally, for ODE models, it supports automatic differentiation and adjoint sensitivity analysis [43, 42]. These features appear ideal for handling tasks like exploring model parameter space, parameter estimation, and automatic model discovery. However, the suitability of Julia for systems biology workflows has yet to be thoroughly evaluated against established tools. For example, even though Julia’s stochastic simulators have been evaluated [32]; they were not compared against performant tools like `RoadRunner` and network-free simulators such as NFSim (available in `PySB`) [52]. Moreover, for ODE models, to date, neither an extensive benchmark has been performed to assess the performance against state-of-the-art packages like `pyPESTO` [49], nor has Julia’s ODE ecosystem been extensively evaluated for common problems arising in biology; like both small and large-scale cellular mechanistic models (e.g. signalling and receptor dynamics), SIR models, and phenomenological models (e.g. spiking). These models encompass various computationally challenging features, such as diverse time scales (e.g. fast and slow chemical reactions), multiple scales (e.g. abundant and scarce proteins), and steady-state criteria (e.g. stationary cells before drug administration).

To evaluate the Julia programming language for workflows with dynamic models in systems biology, we developed `SBMLImporter.jl,` a Julia SBML importer, and `PEtab.jl,` an importer for parameter estimation problems in the PEtab format [50] (Fig. 1a, c). Since many modelling workflows rely on model simulations (Fig. 1), we first used these tools to evaluate Julia’s stochastic simulators (e.g. Gillespie methods) against `PySB` and RoadRunner and deterministic simulators (ODE solvers) against the Sundials ODE suite [19]. Next, since ODE model workflows such as Bayesian inference, parameter estimation, and scientific machine learning often benefit from model derivatives (Fig. 1b-c), we evaluated differentiation methods. Lastly, as a case study, we evaluated parameter estimation performance for ODEs. For the latter two tasks, we compared our results against `pyPESTO,` which utilizes the AMICI interface to Sundial’s ODE suite [19, 11]. We performed all evaluations for ODE models with problems from the PEtab benchmark collection [18, 50], which all include experimental data.

**Figure 1:**
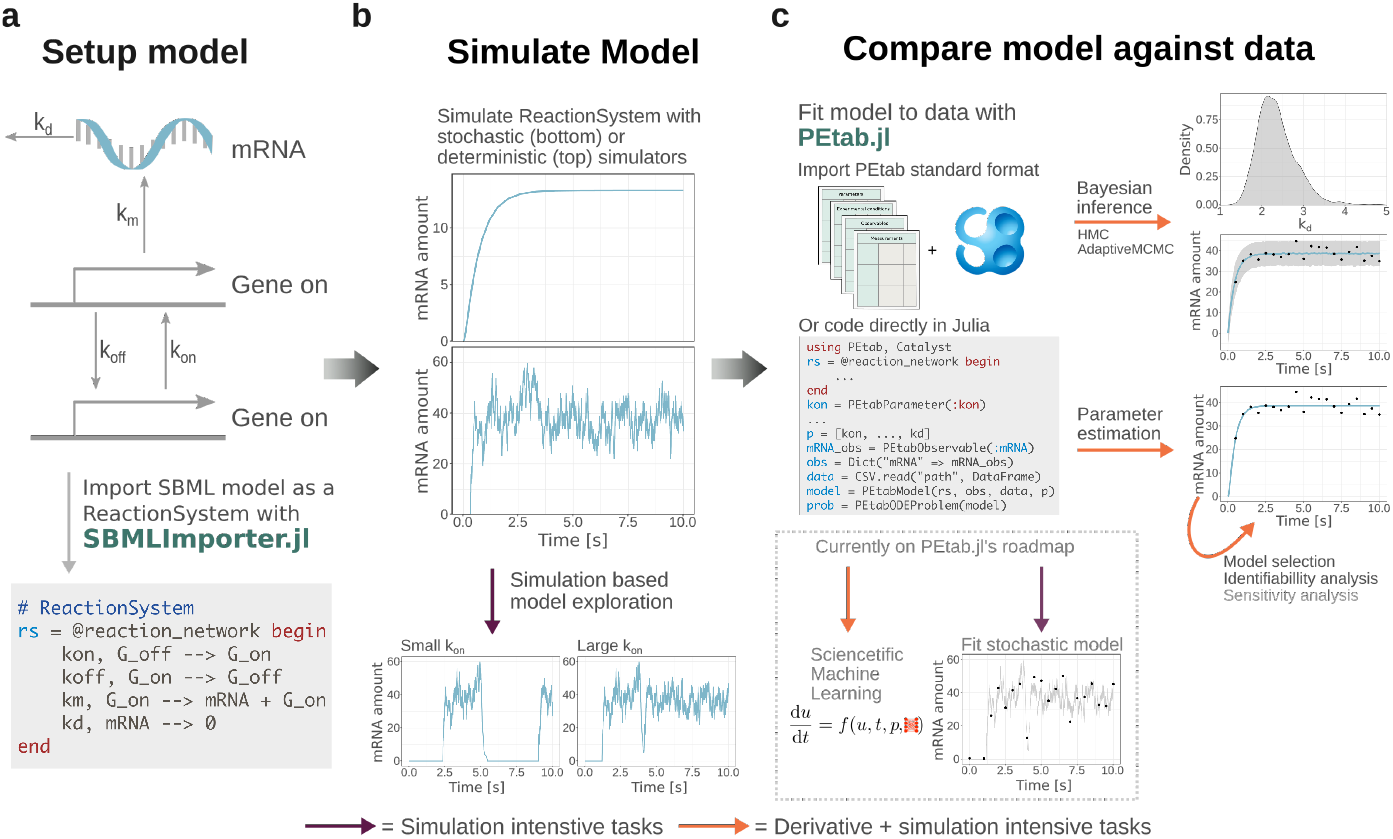
Common modeling workflows and how PEtab.jl and SBMLImporter.jl fit into these. The purple arrows denote tasks that are simulation intensive, and the orange arrows denote tasks that often require both many model simulations and derivative computations. **a)** For a model structure (top) in the SBML format `SBMLImporter.jl` imports the model into a Catalyst reaction network. Note, reaction networks can also be coded directly in Julia (bottom part) **b)** Catalyst reaction networks can be simulated with stochastic simulators or deterministic ODE solvers when for tasks like exploring model behaviour (bottom). These networks are also compatible with additional modelling tools such as `BifurcationKit.jl` **c)** `PEtab.jl` facilitates model fitting by linking measurement data to a model via building a likelihood or given priors, a posterior function. Models in the PEtab standard format can be imported directly, or the model can be coded in Julia, where the dynamic model is provided as a Catalyst reaction system. A parsed PEtab problem can be fitted to data by estimating the unknown parameters (parameter estimation), or the posterior can be inferred by performing Bayesian inference with methods such as adaptive MCMC or the Hamiltonian-Monte-Carlo NUTS sampler (right part). For all these operations `PEtab.jl` supports several methods for computing the gradient of the likelihood/posterior. Currently, `PEtab.jl` does not support scientific machine learning models and inference for stochastic models (bottom), but it is on the road map.

In summary, our tools make Julia more accessible within the systems biology community. Moreover, our quantitative analysis provides guidelines on when to use Julia for dynamic modelling, specifically for choosing a simulator, gradient method, and optimizer.

## Results

### SBMLImporter.jl and PEtab.jl facilitate dynamic modeling in Julia

To make Julia’s ecosystem accessible for models in the SBML standard format, we developed `SBMLImporter.` This package allows users to build models with a wide range of features, such as events and multiple compartments (e.g. cytosol and nucleus), in SBML exportable tools like the Copasi graphical interface [22]. Furthermore, to simplify model fitting workflows, we developed `PEtab.jl` (Fig. 1). Following the `PEtab` standard [50], this package allows users to fit models to data for a wide range of scenarios.

`SBMLImporter.jl` imports models into Catalyst reaction networks [32] (Fig. 1a). This enables stochastic simulations to be converted into a jump-problem (simulated using, e.g. Gillespie algorithm) or a Langevin SDE problem [15]. Alternatively, deterministic simulations can be converted to an ODE problem (Fig. 1b). For each problem, the solvers in the `DifferentialEquations.jl` package are supported [43]. Additionally, Catalyst integrates with additional modeling packages, like `Bifurcationkit.jl` [58]. Note that reaction networks can also be coded directly in Julia. `SBMLImporter.jl` supports SBML features on a similar level of support as established tools like AMICI and `RoadRunner` (Tab. S2).

`PEtab.jl` links ODE models to measurement data by creating a likelihood function or given priors, a posterior function (Fig. 1c). The package handles a wide range of scenarios like different measurement noise formulas for data gathered with different assays, steady-state simulations to capture interventions like drug administration to stationary systems, and data gathered under multiple experimental conditions. PEtab problems in the standard format [50] can be directly imported. Alternatively, problems can be coded in Julia, where the dynamic model is provided as a Catalyst reaction network (Fig. 1c). For the model likelihood, `PEtab.jl` supports both forward and backward gradient computation approaches, suitable for small and large models respectively [7]. Additionally, it integrates with several numerical optimisation packages for parameter estimation and has Bayesian inference support for state-of-the-art methods like Hamiltonian Monte-Carlo [21] and adaptive parallel tempering [54].

`PEtab.jl` and `SBMLImporter.jl` are available on GitHub (https://github.com/cvijoviclab/SBMLImporter.jl and https://github.com/cvijoviclab/PEtab.jl). Both packages can be installed via the Julia package manager. We also provide Jupyter Notebooks and a Google Colab for getting started and/or for performing benchmarks (https://github.com/cvijoviclab/SysBioNotebooks).

### Julia stochastic simulators are often most efficient

Stochastic models in biology can rarely be solved analytically, requiring numerical methods for model simulation. For a given parameter set, stochastic simulators randomly simulate system reactions over time to produce random trajectories [31] (Fig. 1b). Therefore, to characterise model behaviour even for a single parameter set, many simulations are needed. A recent study demonstrated that the simulators in Julia’s `JumpProcesses.jl` are faster than those in BioNetGen, Matlab SimBiology, Copasi, and GillesPy [32]. However, this study did not consider `RoadRunner` and `PySB` [33, 60]. RoadRunner is the simulation engine in several modeling packages such as Tellurium, while `PySB` in particular supports the network-free NFsim method, which was designed for simulating large chemical reaction networks [52]. To benchmark these tools, using `SBMLImporter.jl` we compared four `JumpProcesses.jl` simulators: the direct SSA (Direct) [14], the sorted direct SSA (SortingDirect) [35], Rejection SSA (RSSA), and Composition-Rejection SSA (RSSACR) methods [55, 56, 57], against `RoadRunner’s` Direct SSA method, and `PySB’s` SortingDirect and NFsim [52] methods. We considered the five differently-sized models used for benchmarking in [16] (Tab. 2) that describe processes such as a multistate signalling (multistate, 18 reactions), three-site phosphorylation (multisite3, 288 reactions), epidermal growth factor receptor signalling (egfr net, 3749 reactions), B-cell receptor signalling (BCR, 24 388 reactions), and IgE receptor signalling (Fc*ϵ*RI *γ*2, 58 276 reactions).

To account for simulation initialization overhead, we simulated the models for several time intervals. `JumpProcesses` was fastest for all models (Fig. 2). For the smallest model (Fig. 2a), the SortingDirect method was noticeably faster than `RoadRunner` (2.6 fold speed up at *t* = 10^5^ Fig. 2a). The `PySB` SSA simulator had a large initial overhead, probably due to network generation, which can be sidestepped by directly calling BNGL for model simulations. However, for long simulation intervals where initial overhead has a minor effect, `PySB’`s SortingDirect was comparable for longer simulation times (JumpProcesses had 1.28 fold speed up at *t* = 10^5^). For the two medium-sized models (Fig. 2b, f), RSSACR performed best and was more than four times faster than the best `PySB` method (multisite3 4.1 and egfr net 28 fold speed up compared to `PySB’` s SortingDirect Fig. 2d-e, f). NFsim could not simulate the large BCR model, likely due to disjoint patterns (molecules of the same species not linked by intermediates). For the largest Fc*ϵ*RI-*γ*2 model, RSSACR was roughly 4.6 times faster than NFsim (Fig. 2e).

**Figure 2:**
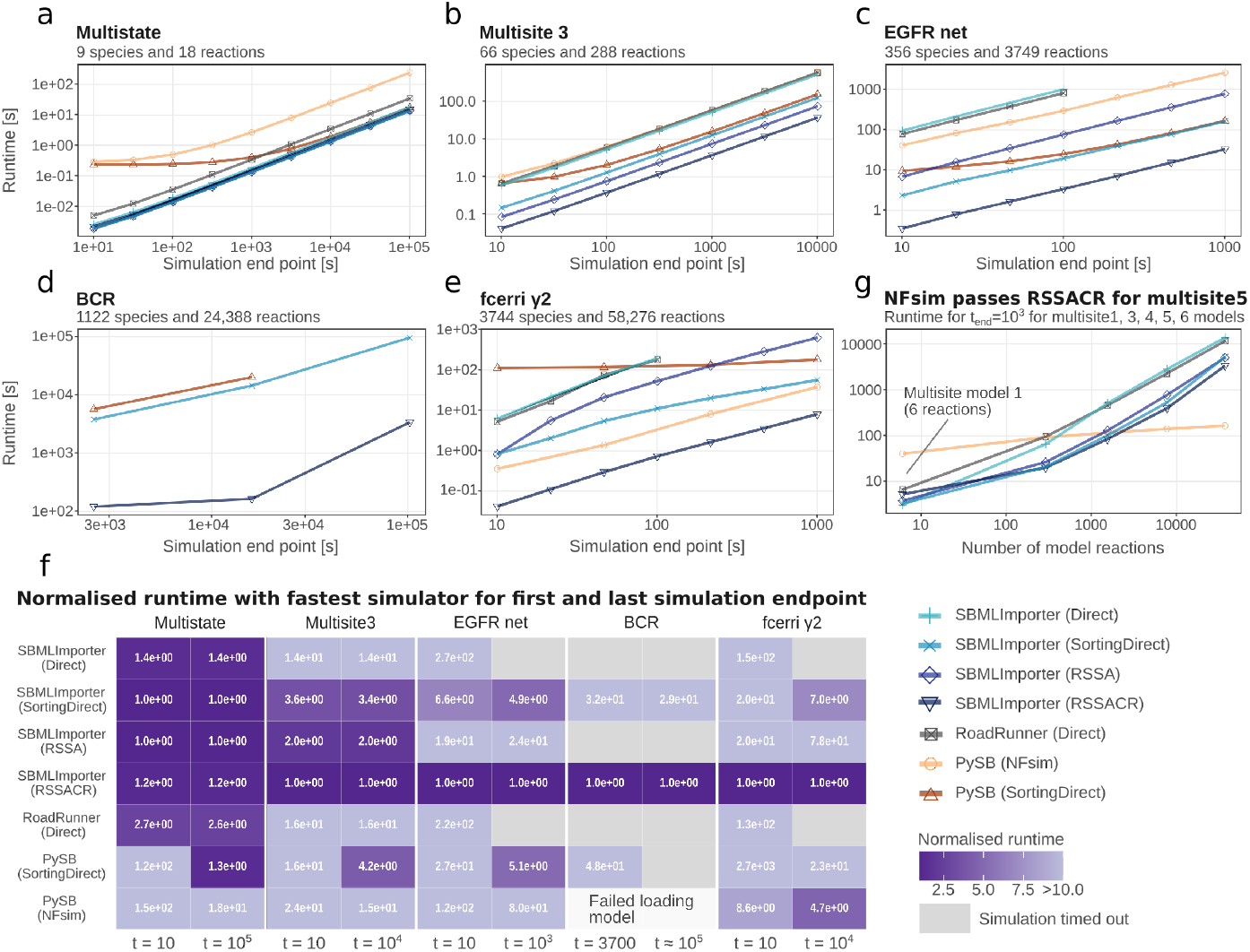
Evaluating stochastic simulators for JumpProcesses (models loaded via SBM-LImporter), RoadRunner and PySB. **a-f)** For `SBMLImporter.jl`, we tested the direct SSA (Direct), sorting direct SSA (SortingDirect), the reaction SSA (RSSA) and composition-reject (RSSACR) methods simulated via `JumaProcesses.jl`. For RoadRunner, the direct SSA method, and for `PySB`, the sorting direct SSA, as well as the network-free NFsim methods. To account for potential initialisation overhead, models were simulated for different endpoints, e.g., the Multistate model was evaluated when simulating to endpoints *t* = 10^1^, …, 10^5^. Evaluated models are **a)** a multistate signalling model, **b)** three-site phosphorylation model, **c)** an epidermal growth factor receptor signalling model, **d)** a B-cell receptor signalling model, and **e)** and IgE receptor signalling model. NFsim failed to simulate the BCR model. `RoadRunner` and SMBLImporter (via JumpProcesses’s Direct and RSSA methods) could simulate the BCR model, but failed to complete the simulations within our imposed 4-day time limit (and are hence not included for this model). **f)** Runtime normalised with the fastest method at the first and last simulation end-point for the benchmarks in panel a-e. For example, for the Multi-state model at *t* = 10^5^ SBMLImporter (via JumpProcesses) SortingDirect was around 2.5 faster than RoadRunner. **g)** Runtime across five multisite phosphorylation models (multisite 1, 3, 4, 5, 6 see Tab. 2) of increasing complexity when simulating each method to a final time of 1000s.

Given that network-free simulators like NFsim often perform well for large models with low molecule counts that have many reactions but few rules (rule-based models), it was unexpected that RSSACR was faster than NFsim for the Fc*ϵ*RI *γ*2 model (has 19 rules and at *t* = 1000 around 6000 molecules). To investigate to what extent RSSACR performs better than NFsim for rule-based models, we tested a set of multisite phosphorylation models with an increasing number of phosphorylation sites where NFsim showed good behaviour [52]. At 5 sites (multsite5, 7680 reactions and 20 rules, Fig 2g) NFsim surpassed RSSACR (2.8 times faster).

Regardless, JumpProcesses’s SortingDirect and RSSACR solvers (loaded via SBMLImporter) performed best overall. Both simulators weakly depend on the order of reactions in the model; for example, to effectively scan the reaction list, the SortingDirect method sorts reactions based on their probabilities. This can be consequential. When importing models into Julia from BioGenNet files via `ReactionNetworkImporters.jl` [23], due to the file format, high-probability reactions often occur early in the reaction list. Therefore, the runtime was, in particular, affected for the Direct and RSSA methods. For example, the Direct was 1.02, 1.12, and 1.70 times faster for multistate, multisite2, and egfr-net models (when loaded using ReactionNetworkImporters), while with SortingDirect sorted direct, the difference was only 0.99, 1.03, and 1.18 (Fig. S1). For the large Fc*ϵ*RI-*γ*2 model, RSSACR’s performance was still notably impacted by reaction order (1.56 times faster with net-file import Fig S1).

In summary, Julia simulators are efficient for small and large models, often being at least four times faster than `PySB` and `RoadRunner` for models with *>* 200 reactions. For large models with low molecule numbers and few rules, network-free simulators can be efficient.

### Optimal ODE solver for model simulations is problem dependent

Most ODE models in systems biology are non-linear and lack analytical solutions requiring numerical ODE solvers to simulate them. The choice of solver algorithm can drastically impact simulation times [43, 53] and even the convergence of the solver [53]. However, given the multitude of available algorithms selecting the best one is non-trivial. To provide guidelines, a recent study extensively evaluated the Sundial’s and LSODA’s solvers for a wide range of models [53] but did not evaluate the ODE solvers available within the Julia DifferentialEquations library [43]. We used `PEtab.jl` to assess model simulation time, reliability (simulation failure frequency), and accuracy of 31 solver algorithms in the `DifferentialEquations.jl` suite, and 2 algorithms from the Sundials suite (Tab. 3). We tested these on 29 benchmark problems varying in size from 3 to 500 states (ODEs), representing a wide spectrum of biological processes such as cellular mechanistic models (e.g. signalling), SIR models (e.g. Covid spread), to phenomenological models (e.g. spiking, cell differentiation) (Tab. 1).

**Table 1.**
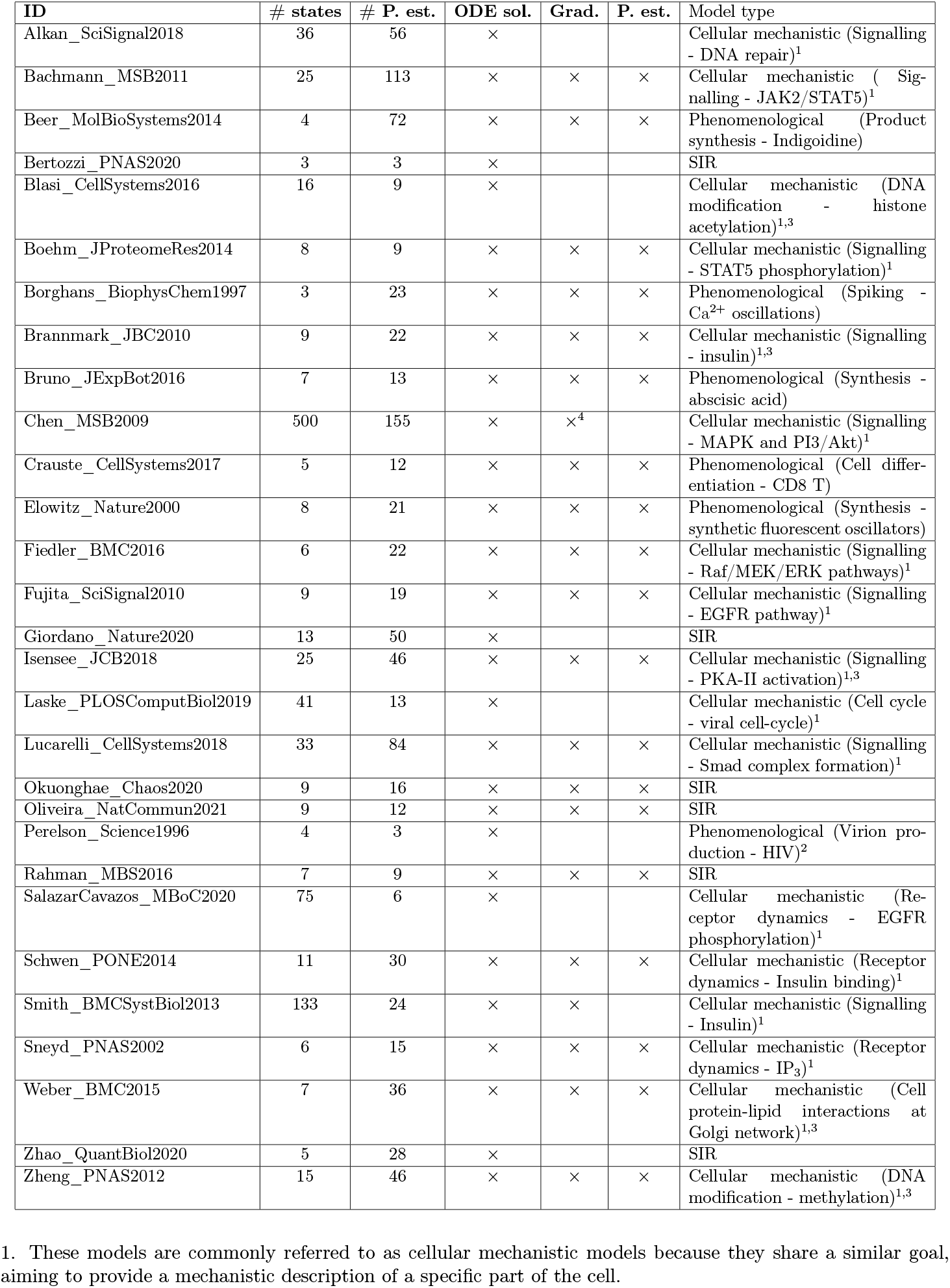
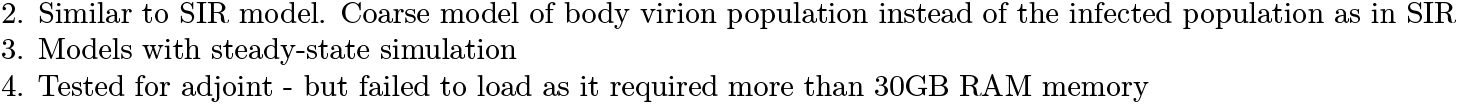
PEtab benchmark problems as specified in the GitHub repository. Denoted are the number of states (ODEs), parameters to estimate, and which specific benchmarks the models were used for (computing the ODE solution, computing the gradient, and performing parameter estimation, respectively). The last column categorizes the model types, including cellular mechanistic models, SIR models and Phenomenological models.

We divided the ODE solvers into four categories, i) Sundial stiff solvers, ii) non-stiff DifferentialEquations solvers, iii) stiff DifferentialEquations solvers, and iv) composite solvers that automatically switch between stiff and non-stiff solvers. Informally, stiffness in ODE models arises when interactions occur on varying time scales, with some being fast (e.g., phosphorylation) and others slow (e.g., translation). Such dynamics are thought to be common in biology [53], and for such models, stiff ODE solvers are beneficial. We found that 11/11 non-stiff solvers failed to simulate 22-31% of the models (blue colour Fig. 3a). Failures occurred primarily for mechanistic models. Notably, non-stiff solvers and composite solvers failed for models with steady-state simulations (4/4 and 2/4 models, respectively), which are commonly used to model interventions like drug administration to stationary systems. Thus, a stiff or composite solver should be used to reduce the risk of simulation failures, which matters during model training, as upon failure, most training algorithms reduce step size, causing longer training times or even the discontinuation of training (and convergence problems). Further, if failures happen near the optimal parameter values, the training process may be biased and never converge.

**Figure 3:**
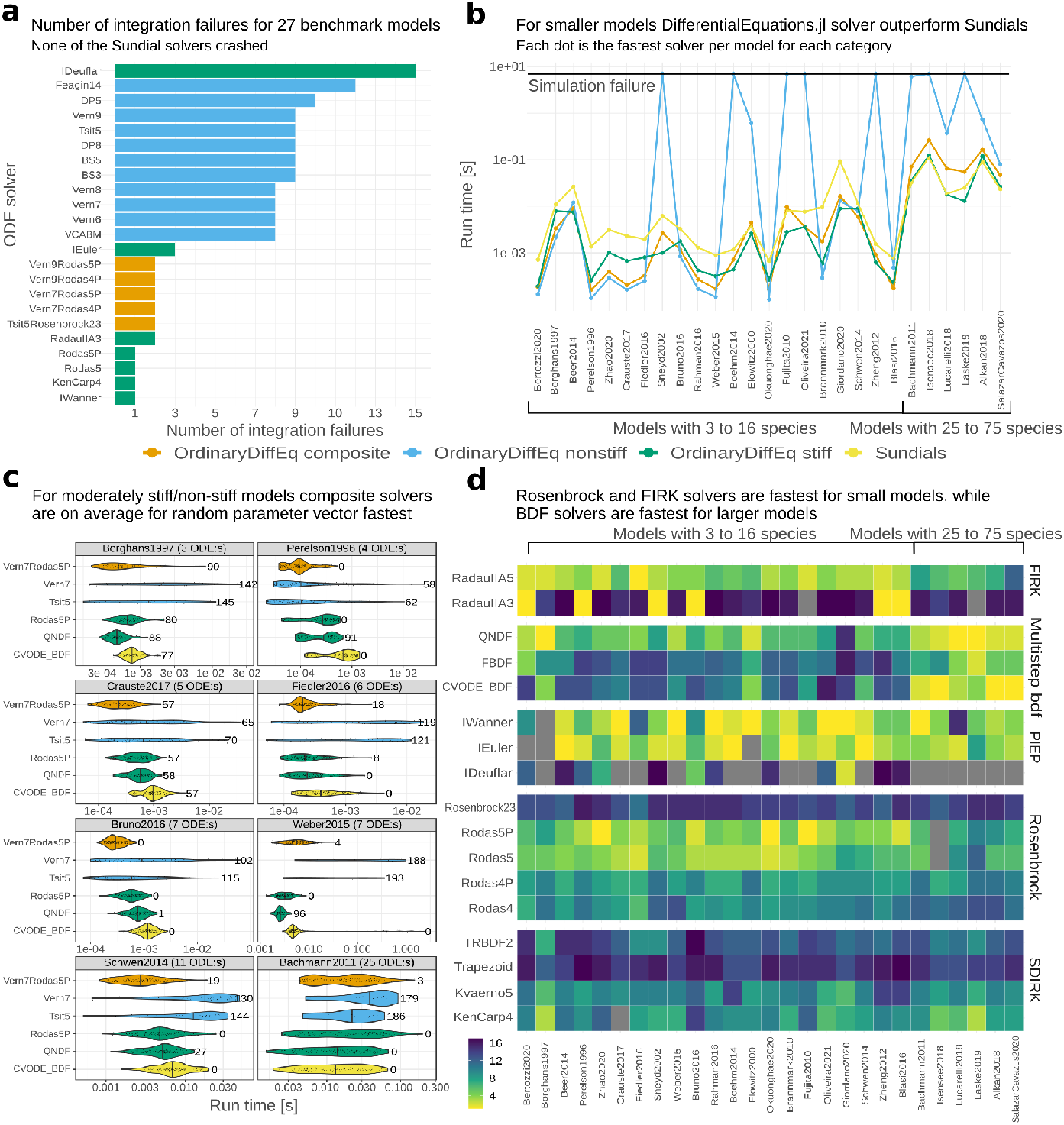
Benchmarking two Sundial and 31 DifferentialEquations ODE solvers for 27 models. Note, that the two bigger models (Chen and Smith) were excluded from this benchmark due to the runtime required to test all solvers. **a)** Total number of integration failures across all models for each solver (Tab. 3). **b)** The best average runtime (averaged over 5 repetitions) for each solver category (Tab. 3). The x-axis is sorted based on the number of states (ODEs) for each model. **c)** Runtime across 200 random parameter vectors for 8 selected models using non-stiff, stiff, and composite solvers that performed well in panel b). The centre line of each violin plot denotes the median, and the number at the end of the violin shows the number of integration failures for that solver, e.g., for the Perelson model, the QNDF failed for 91 out of 200 parameter vectors. **d)** Runtime ranking per model for a large selection of stiff solvers grouped by their respective solver family. A bright yellow colour denotes a good ranking, while a grey colour indicates integration failure.

In addition to reliability, simulation time is important as model training often requires up to 10^3^ − 10^4^ model simulations [18]. `DifferentialEquations.jl` algorithms were fastest for smaller models with up to 16 states (avg. 4.0 fold speedup of Julia stiff (blue) solvers vs Sundials (yellow)), and DifferentialEquations and Sundials were mostly tied for most medium-sized models of 20-75 states (Fig. 3b, avg. 1.2 fold speedup of Julia stiff solvers vs Sundials). Consistent results hold for stricter tolerances (Fig. S2d-e). Noticeably, for ten models (2 mechanistic, 4 SIR, and 4 phenomenological) with ≤ 9 ODEs, non-stiff solvers were the fastest. We suspect this is because the benchmarks were performed at the reported parameter values for which the ODE solution might be well-behaved.

To simulate a real-world scenario where optimal parameters are unknown, we measured the runtime for 200 random parameter vectors (Fig. 3c, Fig. S2c, Fig. S3). For the two mechanistic models (Fiedler and Weber) stiff solvers performed best (yellow and green violins). For phenomenological models (Borghans, Bruno, Crauste, Perlson), composite or stiff performed best (yellow, green and orange violins) and for SIR models, composite (2/4) and non-stiff (2/4) performed best (Fig. S2c). We further looked at two models (Schwen, Bachmann) where stiff/-composite solvers performed best at the reported values, and consistently stiff solvers scored best (yellow and green violins Fig. 3c), that is had fewer simulation failures while being efficient. To further investigate simulation failures, we inspected ODE solver error codes (Fig. S3). Noticeably, for the numerically most challenging models (Crauste and Borghans), composite solvers can get stuck (take *>* 15 min), and while stiff solvers are better, there is no clear best choice for these challenging models (extended discussion in Text S1).

Given the strong performance of stiff solvers, we next looked closer at solver families. Albeit solving the same problem, stiff solvers can be divided into families such as multi-step BDF methods and single-step Rosenbrock methods, each tailored to specific problem types. We found, in line with previous DifferentialEquations benchmarks [43], that Rosenbrock solvers are among the fastest and most accurate solvers for small models (≤ 16 states) (Fig. 3d and Fig. S2b), while for medium-sized mechanistic models (25-75 states) multi-step BDF-solvers are among the fastest. Clear exceptions to this trend are the small Borghans spiking Ca^2+^ model (CVODE-BDF performed best Fig. 3c), and the small SIR Giordinni model (composite solvers performed best Fig. S2c).

For larger models, the performance of stiff solvers depends on the efficiency of the employed linear solver. This is because, at minimum, one linear system must be solved at each time step, and with a dense Jacobian, this scales at least with *(number of ODEs)*2.376 [9]. Julia offers different direct (FastLU, RFLUF, Lapack, Dense, KLU) and implicit (GMRES) solvers, some of which can exploit sparsity in the ODE. For up to medium-sized models (≤ 75 states), the default automatically chosen linear solver for both CVODE-BDF and several DifferentialEquations solvers performed well (dark orange Fig. 4a-c).

**Figure 4:**
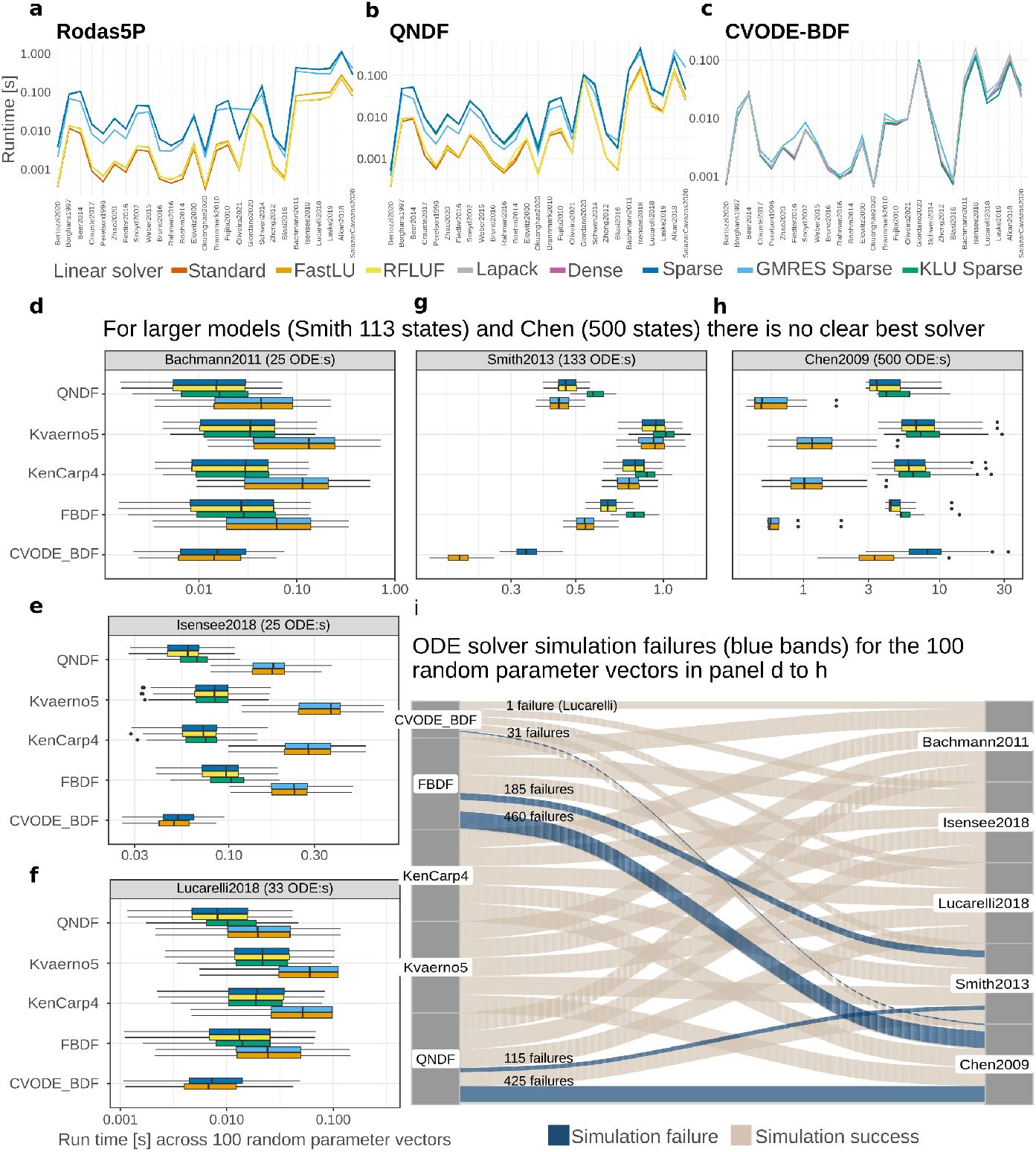
Comparison of different linear solvers for stiff ODE-solvers. **a-c)** Runtime for the Sundial CVODE-BDF and `DifferentialEquations.jl` Rodas5P and QNDF solvers on 27 benchmark models, sorted by size (3-75 states). The colours indicate the linear solver used in the stiff solver time-stepping, with “standard” being the default option and “sparse” indicating the use of a sparse Jacobian. Providing a sparse Jacobian for Julia solvers reduces their performance. **d-h**) Runtime across 100 random parameter vectors for three medium-sized ODE models, and the third (Smith) and second largest (Chen) largest ODE models in the benchmark collection (the biggest requires special infrastructure to work with). **i)** Integration failures are denoted by the blue bands for the 100 random parameter vectors and ODE-solver options in panel d-h, and the number next to the band is the total number of failures for all linear solver combinations for a solver. For example, regardless of linear solver the Julia KenCarp4 solver failed zero times for the Chen model, while the Julia FBDF solver failed for most cases. Results were consistent for different linear solvers, as example, for the FBDF solver, regardless of linear solver the ODE solver ran into integration failure for 92/100 vectors (92 = 460/5). Only for CVODE-BDF did the standard linear solver fail, while the other (KLU sparse) method succeeded.

To study the impact of linear solvers for larger models in a realistic setting for 100 random parameter vectors, we looked at three randomly selected medium-sized models and two newly introduced large models of 113 (Smith) and 500 (Chen) states. For the medium-sized models, BDF solvers (e.g. QNDF and CVODE-BDF) at the default setting performed best (Fig. 4d-h). Meanwhile, for the Smith model, CVODE-BDF combined with a stiff KLU-solver (green violin) was almost twice as fast as the best Julia solvers, while for Chen, the SDIRK solvers (KenCarp4 and Kvaerno5) with a KLU linear solver were more than twice as fast as CVODE-BDF. For both bigger models, Julia BDF-solvers (QNDF and FBDF) frequently failed (Fig. 4i and Fig. S4). The strong performance of the KLU solver might be because it was developed for circuit simulations [8, 9], and circuits might have a topology similar to extensive cellular networks.

In summary, Rosenbrock solvers perform well for smaller mechanistic models (e.g., signalling) and BDF methods for medium-sized models. For large network models, BDF and SDIRK solvers should be compared. Composite solvers scored well for SIR models and a subset of phenomenological models (e.g., cell differentiation). This result indicates that the choice of the best solver is problem-dependent.

### Automatic differentiation accelerates model derivative computations

Common modelling tasks such as model training to find optimal model parameters, identifiability analysis to quantify uncertainty in model predictions/parameters, and sensitivity analysis to identify drug targets typically benefit from accurate gradients [44]. However, for these tasks, gradient computations make out a substantial portion of the runtime (e.g. for the Bachmann model, loss and gradient runtime are 0.02 and 1.1 seconds, respectively). Traditionally for small ODE models, the gradient is computed using forward sensitivities by solving an expanded ODE system [45], where the runtime scales with the size of the expanded system; number of original ODEs (*n*) times the number of parameters (*p*) 𝒪 (*n* × *p*). For larger ODE models, adjoint sensitivity is preferred as the runtime scales approximately as 𝒪(*n* + *p*), with a higher initial cost compared to forward approaches [7, 34]. In contrast to traditional approaches, `PEtab.jl` can utilize forward-mode automatic differentiation (AD) for small models and reverse-mode AD to efficiently compute vector Jacobian products (VJPs) in the adjoint sensitivity analysis computations [34]. To assess whether AD-assisted gradients can accelerate derivative computations and subsequently speed up common modelling tasks, we compared `PEtab.jl` against AMICI [11] -an interface to the Sundials suite.

To compare `PEtab.jl` with AMICI for up to medium-sized models, we selected 18 benchmark models that represent a wide range of biological processes (12 cellular mechanistic, 3 SIR and 3 phenomenological; Tab. 1). We used the CVODE-BDF solver for AMICI, while for `PEtab.jl`, we employed the Rodas5P and QNDF solvers based on previous benchmarks (Fig. 3). First, we focused on the runtime of the loss function. For the 8 smaller models (≤ 20 parameters), there were small differences (AMICI median speedup of 0.9 against both QNDF and Rodas5P), meanwhile, for larger models, AMICI had an advantage (median speedup of 2.1 and 2.6 against QNDF and Rodas5P respectively; Fig. 5a, Tab. S3).

**Figure 5:**
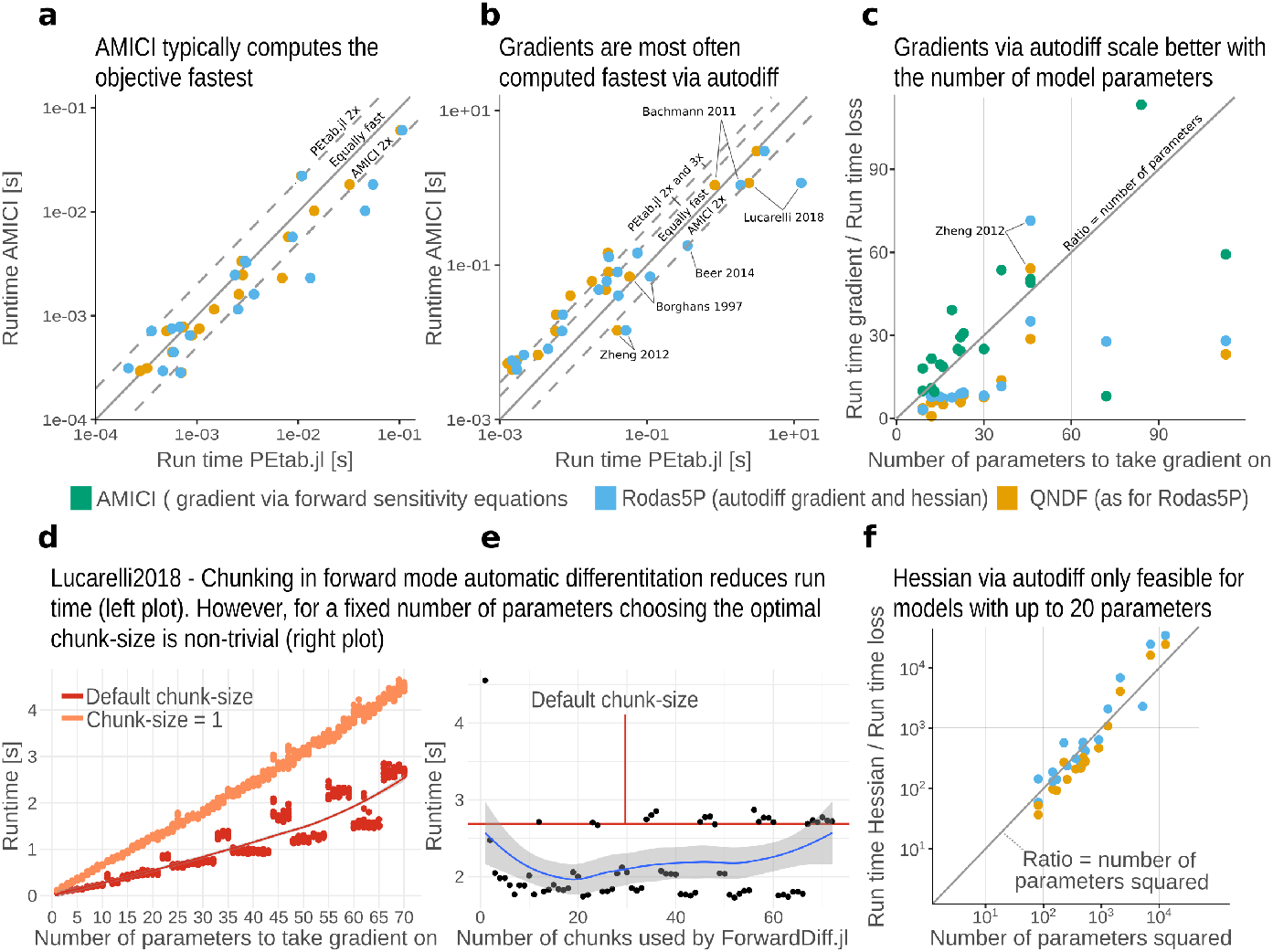
Objective function and gradient benchmark for smaller models. **a)** Runtime evaluation for the loss (objective function) at the reported parameter values. AMICI uses CVODE-BDF, while the Julia PEtab importer uses the Rodas5P and QNDF ODE solvers. Numerical values for panels a-c are in Tab. S3 **b)** Runtime evaluation for the gradient at reported parameter values. AMICI computes the gradient by solving for the sensitivities via an expanded ODE system, while the Julia importer computes the gradient via forward-mode automatic differentiation. **c)** Ratio between the runtime for the gradient in b) and loss in a). The number of model parameters is given on the x-axis. **d)** Gradient runtime for the Lucarelli model when increasing the number of parameters using forward-mode AD without chunking and with the default number of chunks. **e)** Gradient runtime for the Lucarelli model when taking the gradient on all parameters and testing all chunk sizes. **f)** Ratio between runtimes for Hessian calculation. The number of model parameters squared is specified on the x-axis. Absolute values are given in Tab. S4.

Next, we evaluated gradient runtimes. Forward-mode automatic differentiation (AD) outperformed the forward sensitivity approach in AMICI for the majority of models (Rodas5P 67% and QNDF 72% models with median 1.9 and 1.6 fold speedup, respectively), regardless of whether the model is mechanistic, SIR, or phenomenological (Fig. 5b). AD gradients also have better scalability, as only for AD gradients the runtime ratio between the loss and gradient was frequently smaller than the number of parameters (28% AMICI, 89% QNDF, and 94% Rodas5P of the models; Fig. 5c, Tab. S4). Similar results hold for a hybrid approach, in which sensitivity equations, rather than the gradient of the loss functions, are computed via forward-mode AD (Fig. S5a). For higher order derivatives, we found that the loss-Hessian runtime ratio was often smaller than the number of parameters squared (72% QNDF and 53% of the models Fig. 5f, Tab S2-S3). As a coarse rule of thumb, we found Hessian computations feasible (≤ 2 seconds) for models with up to 20 parameters.

As the efficiency of the forward-mode AD methods was surprising, we next analysed AD tuneable options to understand why. In a naive AD implementation, a single directional derivative is computed per forward pass, thus as many passes as parameters are needed to compute the gradient [1], resulting in a linear scaling. `ForwardDiff.jl` employs a technique called chunking, which allows for a tunable number of directional derivatives to be computed per forward pass [46]. We tested the runtimes of chunked and un-chunked AD for three models where we successively increased the number of parameters on which to compute the gradient. Chunked gradients scaled better (Fig. 5d and Fig. S5), but it should be noted that chunk size is a tuning parameter, and the default value may not be optimal (Fig. 5e and Fig. S5). Why chunking improves performance is unclear, but at least for simple cases, compiler output suggests that chunked gradient operations frequently use efficient vectorised CPU SIMD instructions. Therefore forward-mode AD might have an advantage over other approaches because it effectively leverages modern CPU architectures.

For large ODE models with more than 100 ODEs + parameters, such as extensive signalling or metabolic networks, forward approaches are impractical (e.g., a single forward-mode AD gradient for the Smith model requires ≈ 100 seconds). Here adjoint sensitivity analysis can drastically reduce runtimes [7, 34], but runtimes are still substantial, so selecting the best adjoint algorithm is important. `PEtab.jl` supports both the interpolation and quadrature algorithms from `SciMLSensitivity.jl`, along with reverse-mode AD approaches to compute the vector Jacobian product (VJP) in the adjoint ODE [42, 34]. To compare these algorithms with AMICI, we selected five representative benchmark models ranging from 17 to 655 parameters+states (1 small, 2 medium-sized, and 2 large) and evaluated gradient runtime at 50 randomly selected parameters. AMICI proved the most reliable (9.5% and 64% average fail rate for AMICI and the best SciMLSensitivity algorithm, respectively; 6b). Furthermore, `PEtab.jl` failed to load the large Chen model (655 parameters + states) as it required ≥ 30GB of RAM. Excluding the failed runs, `PEtab.jl` was faster (avg. speedup of 1.3 and 2.1 for the Boehm and Bachmann models respectively; Fig. 6a, c-d), and was often more accurate (Fig. 6c-d) when we used the interpolation approach and the Enzyme automatic differentiation library to compute the VJP [39]. This is likely because the Enzyme automatic differentiation approach to compute the VJP in the adjoint ODE is more efficient than computing it directly via sparse-matrix representations as done in AMICI.

**Figure 6:**
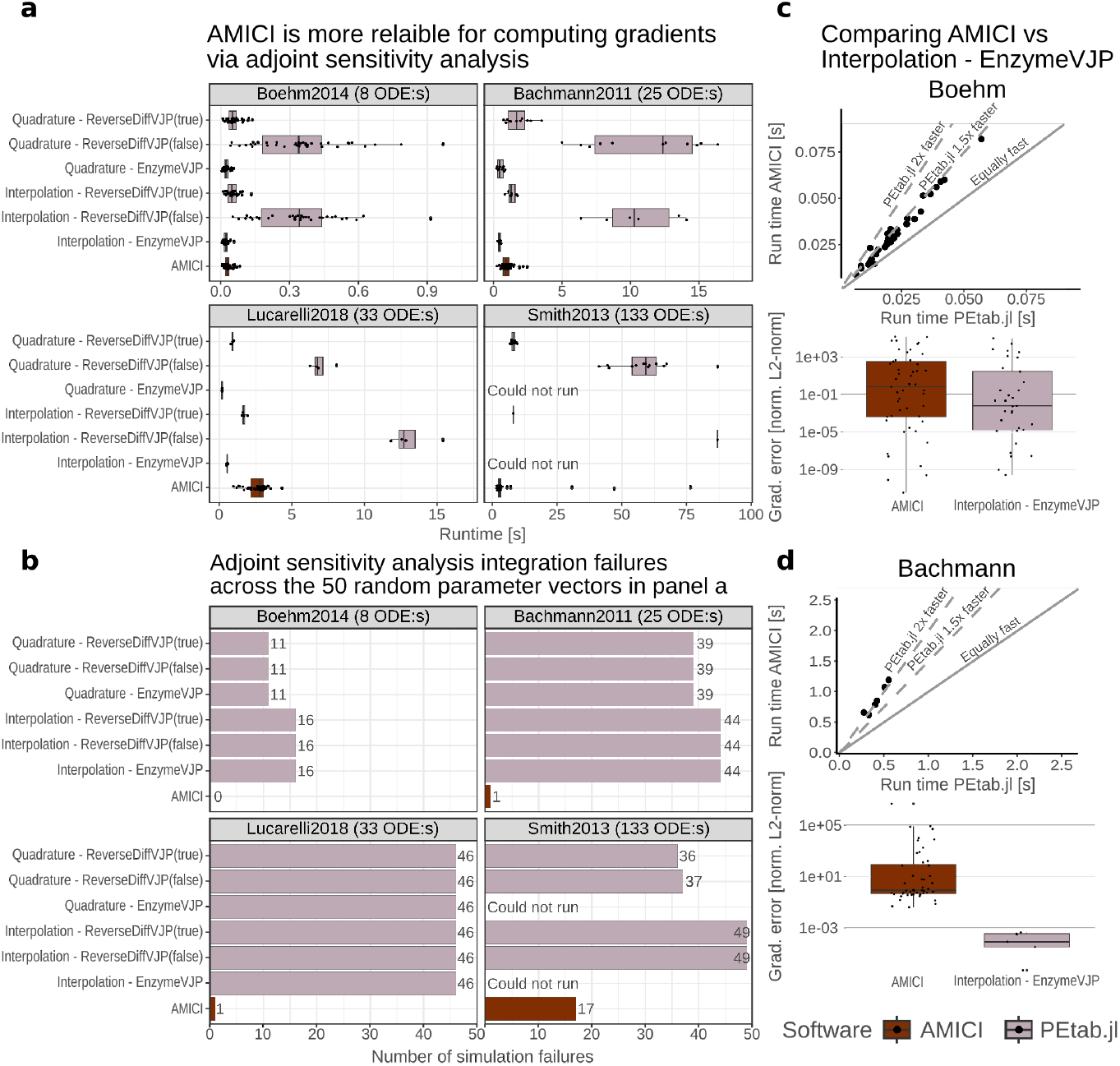
Benchmarking adjoint sensitivity gradient computation methods for five larger ODE models. **a)** Runtime across 50 random parameter vectors when computing the gradient via adjoint sensitivity analysis. We tested two approaches for `PEtab.jl` : the quadrature and interpolation approach in `SciMLSensitivity.jl`, with different ways of computing the vector-Jacobian-product in the adjoint ODE. For instance, “ReverseDiffVJP(true)” refers to using `ReverseDiff.jl` with a compiled tape (true). For Julia, we used the CVODE-BDF solvers as they performed best. **b)** Integration/gradient failures when solving the adjoint ODE for the 50 random parameter vectors and ODE-solver options in panel a). Notably, the AMICI approach, which employs the adjoint algorithm in Sundial, is by far the most reliable method, as for the bigger models (e.g. Smith), all methods in `PEtab.jl` frequently failed. **c**,**d)** Runtime (upper) and accuracy (lower) comparison between AMICI and the interpolation approach in SciMLSensitivity using the Enzyme automatic differentiation library to compute the VJP for the Boehm and Bachmann models. When SciMLSensitivity does not fail, it consistently outperforms AMICI. Gradients were computed by ∥∇*f*_*adjoint*_ − ∇*f*_*ref*_ ∥_2_ where the reference gradient ∇*f*_*ref*_ was computed using a high-order finite-difference method.

In summary, for smaller models (≤ 75 ODE + parameters), whether they are mechanistic, SIR, or phenomenological, forward-mode automatic differentiation is generally faster than the forward sensitivity analysis approach for gradient computations. For larger mechanistic models, the adjoint algorithms from SciMLSensitivity are fast but currently less reliable than the adjoint sensitivity analysis approach accessible via AMICI based on CVODES.

### PEtab.jl often facilitates training efficiency of models

ODE-based models in systems biology often have unknown parameters, such as reaction rate constants, that must be estimated from experimental data. This training problem corresponds to a continuous optimization problem, and to accurately evaluate the quality of a model to, for example, compare it against other models (hypotheses), it is crucial to identify the best fit, preferably rapidly. To evaluate `PEtab.jl` we set up a benchmark against `pyPESTO` which leverages AMICI for model simulations using the same benchmark models as for the gradient evaluation (12 cellular mechanistic, 3 SIR and 3 phenomenological; Tab. 1). For `pyPESTO`, we used the Newton-trust region Fides training algorithm [10] and tested three different approaches to approximate the Hessian: default, BFGS, and Gauss-Newton. For `PEtab.jl`, we tested Fides with the BFGS and Gauss-Newton (GN) Hessian approximation, along with the `Optim.jl` [38] Interior-point Newton (IPNewton) method with GN hessian approximation. Moreover, since `PEtab.jl` can compute the full Hessian for models with a Hessian runtime ≤ 2 seconds, we tested Fides and IPNewton with a full Hessian. For each model, we performed 1000 optimization runs using Latin-Hypercube-generated random start guesses. Model-specific convergence plots can be found in the supplementary (Fig. S8-S26).

Two important model training evaluation criteria, exemplified by the Fiedler model (Fig. 7a-c), are overall runtime and the number of runs that converged to the best likelihood value. For the Fiedler model, the Interior Point Newton method (IPNewton, yellow) had a higher convergence rate compared to Fides BFGS (grey) (5.8% and 0.2% respectively). However, BFGS was faster (IPNewton: 66.6h; BFGS: 1.51h). A summary of the overall runtime and the number of converged runs is the overall efficiency (OE)-score, which is the number of converged runs divided by the runtime [59]. The OE-score provides the number of converged starts, which can be expected per unit of time, meaning that larger values are better. For the Fiedler mode, the BFGS achieved the highest and thus best score (BFGS 1.32OE and IPNewton 0.87OE).

**Figure 7:**
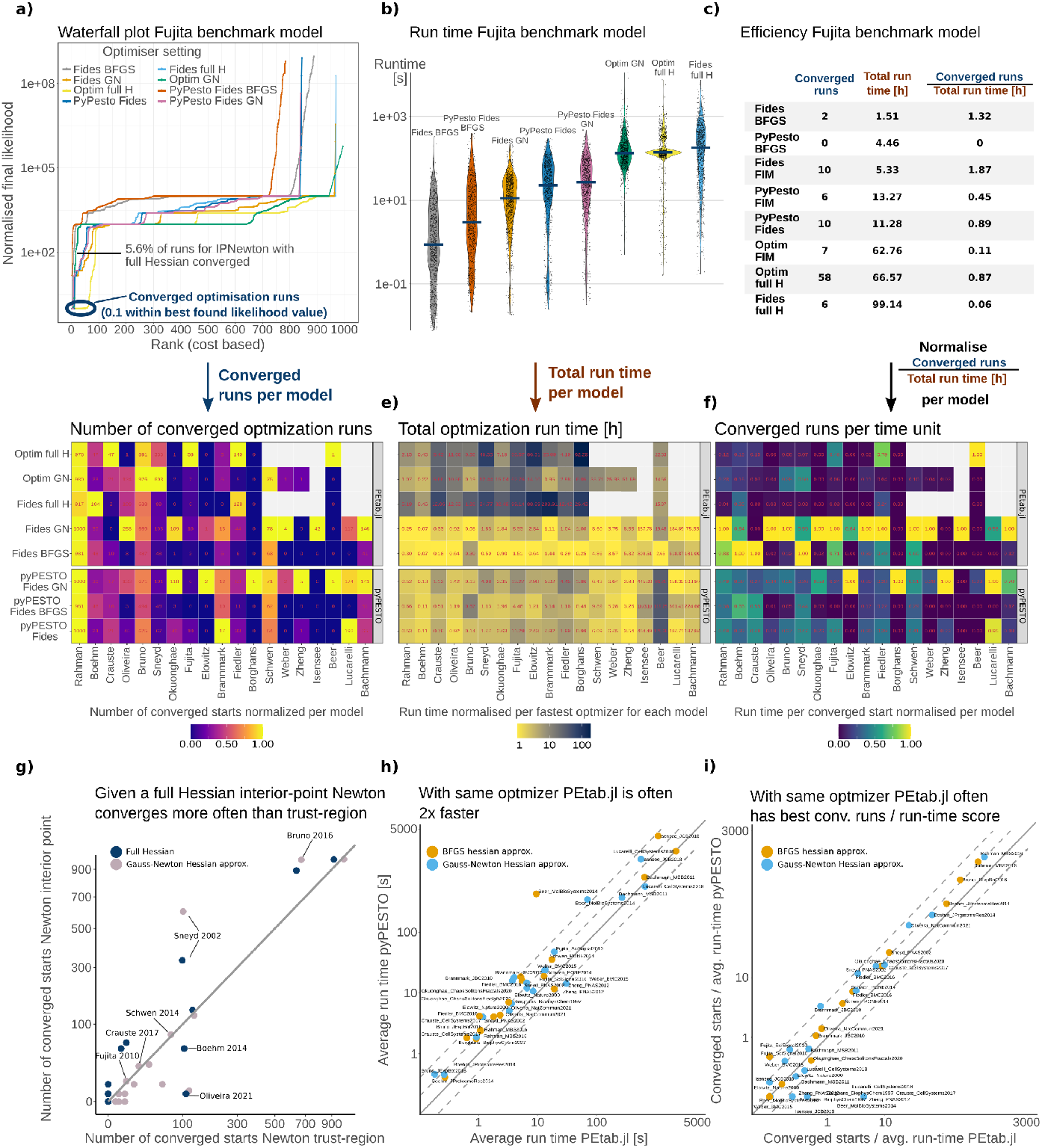
Parameter estimation benchmark for pyPESTO against PEtab.jl. **a)** Waterfall plot for the Fujita model from 1000 optimisation runs. The trust-region Newton optimizer Fides and the interior-point Newton method Optim were used, and the second word indicates the Hessian computation method, such as full Hessian refers to computing the full Hessian via forward mode AD **b)** Runtime for the runs in a) for the Fujita model. **c)** Parameter estimation statistics for the Fujita model, including the efficiency measure of the number of converged runs divided by runtime. **d)** Number of converged runs from running 1000 optimization runs for each model (x-axis, sorted by the number of parameters) and optimizer option (y-axis) from random starting guesses. The color indicates the number of converged runs normalized by the best method per model. White boxes denote that the method was too expansive to run. **e)** Total wall time in hours to perform the 1000 optimization runs for each model and optimizer. The color indicates runtime normalized with the fastest option per model. **f)** Converged run per time unit (panel a divided by panel b) normalized per model, where matches the values in the heatmap. **g)** Number of converged runs for the interior-point Newton method in `Optim.jl` and the Fides trust region method. The dark blue indicates the full Hessian was computed, and in this setting, the interior-point method outperformed Fides. **h)** Average runtime for `pyPESTO` and `PEtab.jl` when using the Fides optimizer. Note that `PEtab.jl` is often more than twice as fast as pyPESTO. **i)** Comparison of the number of converged runs divided by runtime when using the Fides optimizer for `pyPESTO` and `PEtab.jl`, as in h).

When considering the OE-score across all models, `PEtab.jl` outperformed `pyPESTO` for 15/19 models (3 SIR, 1 phenomenological and 11 cellular mechanistic models), with `pyPESTO` performing better for the Borghans, Elowitz, Lucarelli and Zheng models. The Zheng model requires a pre-equilibrium (steady-state simulation), for which AMICI has sophisticated handling. For the Lucarelli model `pyPESTO` had a notably higher number of converged starts when using the Gauss-Newton Hessian approximation, but for the Oliveira and Isensee models `PEtab.jl` had more converged runs. Looking closer at two models with differences in converged runs (Crauste and Isensee), computing the Gauss-Newton approximations proved challenging (even with low 10^*−*8^ ODE solver tolerances), implying that the impact of using different ODE solvers and gradient methods likely is what affected convergence (Text S2, Tab. S1). Lastly, for Elowitz, Borghans and Beer, neither software frequently converged, so results should be interpreted carefully. The low convergence rates are likely related to model characteristics, both Elowitz and Borghans are stiff and oscillate which is hard to handle [40]. Beer has many parameters to estimate (72) where a subset corresponds to the activation time of an event, which require special methods [6]. Regardless, training options where we used a Hessian approximation instead of the full Hessian consistently achieved the best score (18/19 models), as computing the latter is computationally demanding.

`PEtab.jl` often achieved the best OE score due to its faster speed. To quantify the speed difference, we compared the runtime when using the Fides optimizer for both software. For all but two models (Zheng, Lucarelli), `PEtab.jl` was faster (avg. speedup 2.31 fold with respect to Fides GN and 4.44 fold with respect to Fides BFGS Fig. 7h).

While providing a full Hessian did not improve the overall OE score, it did improve convergence for some types of models (for 0/3 SIR, 1/3 phenomenological, 5/7 mechanistic models). For instance, in the Fiedler model, the best Hessian approximation converged in only 1.0% of runs (Fig. 7a-c), whereas the IPNewton method with full Hessian converged in 5.8% of runs. However, providing the full Hessian does not always yield better results, as seen for the Brannmark, Bruno, Borghans, Elowtiz, Okuonghae, Oliveira and Rahman models.

Lastly, we evaluated the earlier claim that trust-region methods often perform best by comparing results from the Newton interior-point and trust-region Newton methods [18]. When we used the Gauss-Newton Hessian approximation, similar to earlier studies [44, 18], the trust-region method was indeed best (for 12/16 tested models). However, when we computed the full Hessian, the interior-point method outperformed the trust-region method (for 11/13 of tested models Fig. 7e). Further, for 7/13 and 2/13, the complete Hessian interior point and trust-region methods, respectively, performed better than the `pyPESTO` trust-region with Gauss-Newton approximation.

In summary, computing a full Hessian can improve training convergence, especially for mechanistic models. However, it comes at a computational cost, which questions its applicability for models with 15 or more parameters. When we can compute the full Hessian, interior-point algorithms often perform better, whereas it is feasible only to compute a Gauss-Newton Hessian approximation, trust-region methods tend to perform better.

## Discussion

Dynamic modeling plays a pivotal role in understanding cellular dynamics, which are essential for our understanding of the functioning of living organisms. And doing it fast enhances the efficiency and utility of this process. Here, we introduced a Julia SBML importer, `SBMLImporter.jl`, and `PEtab.jl` -a package for fitting models to data based on the PEtab format [50]. Using `SBMLImporter.jl`, we evaluated stochastic simulators against `RoadRunner` and `PySB`, and in general, Julia solvers were the fastest. For ODE models, using benchmark models with real data covering a wide spectrum of biological processes such as cellular mechanistic models, SIR models, to phenomenological models capturing core mechanisms, we evaluated common modelling workflows like model simulations, computing model derivatives, and estimating unknown parameters from experimental data. For the latter two workflows, `PEtab.jl` was frequently more than twice as fast as the state-of-the-art `pyPESTO` package for models with ≤ 75 parameters. However, for larger mechanistic models, derivative computations using adjoint sensitivity analysis were less reliable.

Based on the model training (parameter estimation) evaluation criteria, converged runs per time unit (OE score), `PEtab.jl` performed better than `pyPESTO` for 15/19 models. `PEtab.jl` consistently achieved the best OE score when a Hessian approximation is computed instead of the full Hessian. However, a full Hessian generally improved convergence for non SIR models (note only 3 SIR models were tested). Thus, at least for non-SIR models, if optimization runs can be performed in parallel on a multi-core computer, computing the full Hessian is likely worthwhile if it takes less than 1 second. Additionally, `PEtab.jl` computes the Hessian via a forward-over-forward approach (quadratic scaling with the number of parameters), but if the challenging engineering task of implementing an adjoint-over-forward approach (linear scaling) can be solved [41], the improved convergence properties can be leveraged at lower run times.

`PEtab.jl` often performed better than `pyPESTO` because it was faster. This is not due to symbolic model pre-processing, as both software do it. Rather, for smaller models `PEtab.jl` has access to ODE solvers tailored for such models (Fig. 3). In addition, `PEtab.jl` computes gradients faster via forward-mode AD than AMICI computes them via forward sensitivity analysis. Consistent with earlier studies [36], we found that chunking, which means computing multiple directional derivatives per forward pass, is the primary reason for the low runtimes of AD gradients. We also noted a typical non-linear relationship between chunk size and gradient computation time, so although tuning it can be beneficial, it is non-trivial. When dealing with larger models that require adjoint sensitivity analysis `PEtab.jl`, which uses the quadrature or interpolation methods from `SciMLSensitivity.jl`, frequently failed to compute the gradient. Meanwhile, AMICI, which uses the well-established Sundials suite, proved more reliable. When `PEtab.jl` did not fail, it was 1.5-2x faster, suggesting that Julia has the potential to speed up training for large models, but further work is needed to make adjoint methods more reliable.

In addition to several gradient methods, `PEtab.jl` has access to the ODE solvers from the `DifferentialEquations.jl` suite. This is beneficial, as Rosenbrock solvers proved effective for small mechanistic models, composite solvers for SIR models, and BDF and SDIRK for larger mechanistic models. For phenomenological models, we did not identify any clear guidelines, as for some models like Crauste (cell differentiation), composite solvers performed the best, while for others like Borghans (spiking), stiff solvers performed best. This might depend on the level of mechanistic details in the models, as models with simplified mechanisms such as SIR models may have states that exhibit similar time scales and, thus, less stiffness suitable to composite solvers. Regardless of model type, contrary to the belief of the prevalence of stiffness in biological models, we found that non-stiff solvers performed well at reported parameter values for 10 out of 29 models. However, except for two SIR models, this did not hold for random parameter vectors. Therefore, for most workflows, a stiff or composite solver should be used.

The results for stiff ODE solvers can also be examined from an algorithmic viewpoint. For small models, the main computation time is spent on evaluating the ODE rather than solving linear systems [9]. Therefore, Rosenbrock solvers, which perform multiple-linear solves per time-step to take long steps, are efficient [48]. For larger models, the runtime is often dominated by linear solves, thus, multi-step BDF solvers like Sundials CVODE-BDF that perform fewer linear solves per time step are efficient [19]. However, multi-step methods require a start-up phase with shorter steps at the start or following an event (e.g., insulin dosage). Hence, for models like Chen with multiple events, single-step solvers such as SDIRK methods [29], designed to minimize linear solve time, can be preferable, as seen in our results (Fig. 4). Overall, this suggests that for models of biological systems, Julia is likely to offer speed improvements compared to suites like Sundials, primarily for smaller models and larger models with numerous events. To guide the users, we provide a flow chart for choosing the ODE solver (Fig. S6, S1 text).

In addition to ODE solvers, we evaluated the stochastic simulators in `JumpProcesses.jl`, which consistently outperformed `PySB` and RoadRunner. Considering previous benchmarks where Julia outperformed BioNetGen, GillesPy, Matlab, and COPASI [32], this cements the efficiency of Julia’s simulators. For larger models (*>* 200 reactions), Julia’s RSSACR method performed best and was even faster than the network-free NFsim method. However, while NFsim cannot simulate certain challenging model structures such as the BCR model Fig. 2d), it outperformed RSSACR for a model with low molecule count, few rules, and many reactions. This highlights that NFsim is favoured for a certain class of models, such as polymerisation and large complex formation models. Nonetheless, our benchmarks align with the `JumpProcesses.jl` documentation guidelines [24] that SortingDirect and Direct are suitable for smaller models, while for larger models RSSACR should be preferred.

Taken together, the results of our extensive benchmark analysis lead to a natural question: when is it suitable to use the Julia programming language for dynamic modelling in biology? For simulation-intensive workflows such as exploring model behaviour across millions of parameter sets, Julia excels. For the stochastic models, the simulators are efficient, and there is only a small subset of scenarios where network-free simulators (NFsim) are preferred. For ODE models, access to a wide selection of ODE solvers for both small and large problems makes Julia suitable for most models. Importing models is also straightforward, with tools supporting both the BioGenNet `.net` format and now extensively the SBML format. For gradient-dependent ODE workflows, Julia is highly performant and recommended for small mechanistic, SIR, or phenomenological models. Thanks to efficient ODE solvers that are compatible with forward-mode AD, `PEtab.jl` was often more than twice as fast as AMICI. Furthermore, seamless integration of AD with ODE solvers offers benefits such as easy Hessian computations for improved training convergence, and as available via `PEtab.jl` efficient Bayesian inference via Hamiltonian Monte Carlo (HMC) samplers [12]. Moreover, our results suggest that, with additional engineering, Julia can accelerate the training process for larger stiff models.

Extending beyond the current scope of `PEtab.jl`, automatic differentiation presents an opportunity to streamline the development of new methods for computationally demanding tasks such as model discovery via scientific machine learning [42] (Fig. 1c). From a broader perspective, Julia is a powerful tool with a large potential for dynamic models in systems biology. Furthermore, the language already has mature packages for distributed computing, facilitating tasks like parallelizing workflows such as model exploration across millions of parameters. However, to fully harness its demonstrated potential, the next step would be the development of tools like `pyPESTO` and `D2D` that support the entire modeling pipeline [45, 49]. This presents an exciting opportunity for the community to collaborate and build upon existing frameworks, paving the way for more efficient and accessible computational biology tools.

## Materials and methods

We evaluated `PEtab.jl` against `pyPESTO` and AMICI on previously published models with real experimental data from the PEtab benchmark collection [18, 50]. A summary of key features and benchmarking information for each model is available in Tab. 1. For testing gradients, we selected a subset of models with a wide range of parameters to estimate (9 - 155), covering several biological features (mechanistic, SIR and phenomenological) and a wide range of computational features such as parameter-dependent initial conditions with different observable function and noise models, preequilibration, and log-transformations. To benchmark parameter estimation performance, we used the same models as for the gradients, except for the larger Smith and Chen models as `PEtab.jl` achieved poor performance when computing gradients for large models via adjoint sensitivity analysis (Fig. 6). We divided the models into three categories based on which process they modelled: (1) cellular mechanistic models, (2) SIR models and (3) phenomenological models (e.g. spiking, virus growth). The classification is not based on trajectories (e.g. some SIR models might have similar ones to phenomenological models), but rather the process they model, e.g. SIR models all focus on disease progression in a population.

For stochastic model simulators, we used the five models previously evaluated in [16, 32], Tab. 2, and for additional testing of NFsim, we used a series of multi-site phosphorylation models with increasing number of binding sites from [52]. These models cover a range of scenarios, like phosphorylation, receptor dynamics, and signalling. Moreover, as different stochastic simulators are often suited to a certain model size, these models cover a wide range of sizes (Tab. 2).

**Table 2.** PEtab benchmark problems for evaluating stochastic model simulators. Listed are the number of model species, reactions, and parameters and since all models are generated from the rule-based BioNetGen language, the number of rules.

| ID | # Species | # Reactions | # Parameters | # Rules |
| --- | --- | --- | --- | --- |
| Multistate | 9 | 18 | 9 | 4 |
| Egfr net | 356 | 3749 | 43 | 23 |
| BCR | 1122 | 24 388 | 128 | 76 |
| Fceri $\gamma 2$ | 3744 | 58 276 | 26 | 19 |
| multisite1 | 6 | 6 | 9 | 4 |
| multisite3<sup>1</sup> | 66 | 288 | 9 | 12 |
| multisite4 | 258 | 1536 | 9 | 16 |
| multisite5 | 1026 | 7680 | 9 | 20 |
| multisite6 | 4098 | 36 864 | 9 | 24 |
| multisite7 | 16 386 | 172 032 | 9 | 28 |

1. We downloaded multisite models from http://michaelsneddon.net/nfsim/pages/models/models.html, where due to a mistake, the code for multisite 2 is actually the one for multisite 3 (hence we did not benchmark multisite 2)

### Stochastic model formulation

Stochastic model simulations in systems biology are often carried out using so-called jump processes. These are continuous-time, discrete-space simulations, simulating the actual reaction events of the system and their impact on its state. How these jump processes are generated from a model is described by stochastic chemical kinetics [31]. In practice, these stochastic chemical kinetics-based jump process simulations are carried out using some algorithm, of which Gillespie’s (or the stochastic simulation) algorithm is the most well-known one [13, 14].

### Implementation in this study

To simulate stochastic chemical kinetics, we use the `JumpProcesses.jl` Julia package [24, 61]. Specifically, we used Gillespie’s direct method, the sorting direct method, and the rejection and composition-rejection SSA methods [14, 35, 55, 56, 57]. We also use the `PySB` and `RoadRunner` simulation packages [33, 60]. We used their implementation of Gillespie’s direct method (`RoadRunner`), the sorting direct method (`PySB`), or NFsim (`PySB`) [52]. All trialled simulators are exact methods. This means that they sample the exact jump process described by stochastic chemical kinetics. Hence, their performance can be directly compared without taking tolerance, error, or similar measures into account.

To ensure only the computational cost of the actual simulations was measured, we disabled all saving of intermediary solution states (i.e., we only saved the solution at the initial and final time points). For `JumpProcesses`, we used the `save_positions = (false, false)` option (which disables solution saving before and after jumps). For PySB, we used the `n_runs = 1` (limiting simulation runs to a single simulation) and `gml = 50000000` (a safety which throws an error when molecule numbers are high, increasing it is required to enable simulation for large models) options. We used the RoadRunner’s Julia binding (with no additional options).

### ODE model formulation

We consider ODE models that can be expressed as:

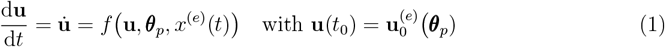

Here, ***θ***_*p*_ are the parameters that govern the system’s dynamics of the system. The solution vector **u** ∈ ℝ^*m*^ has *m* components and depends on a condition-specific input function *x*^(*e*)^(*t*) and an initial value function 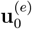. Note that the initial values can depend on the parameters themselves.

#### Implementation in this study

To solve the ODE models `PEtab.jl` uses the Julia `DifferentialEquations.jl` or `Sundials.jl` packages [20, 43]. For the gradient and parameter estimation benchmarks, we used the solver tolerances *abstol* = *reltol* = 1 × 10^8^ for both AMICI and our PEtab importer. For the ODE solver benchmark (Fig. 3), the tested solvers are reported in Tab. 3.

**Table 3:**
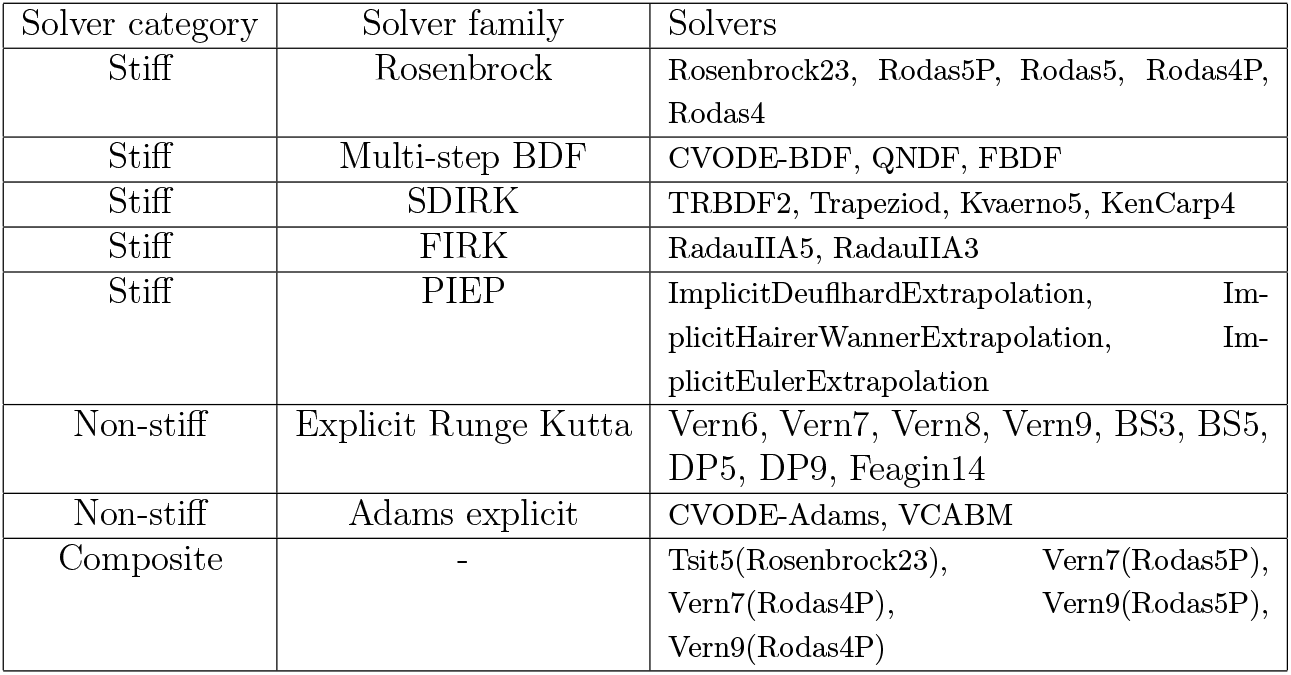
ODE solvers evaluated. Each ODE solver is divided into a solver category and solver family. While each solver solves the same problem, each solver family has different properties, e.g. Rosenbrock is single-step implicit, while mult-step BDF solvers are multi-step methods. Families are Rosenbrock [48], multi-step backward differentiation formula methods (BDF) [20], singly-diagonal implicit Runge-Kutta methods (SDIRK) [29], Fully-Implicit Runge-Kutta Methods (FIRK) [17], Parallelized Implicit Extrapolation Methods (PIEP) [5], Adams explicit [20], and explicit Runge Kutta. For the composite solvers, the first name denotes the non-stiff solver, and the second names the stiff solver that the composite solver should switch to upon stiffness detection.

### Parameter estimation for ODE models - problem formulation

We define ***θ*** = (***θ***_*p*_, ***θ***_*n*_) as a set of parameters, where ***θ***_*n*_ are parameters not a part of the ODE system and ***θ***_*p*_ are a part of the ODE system (or initial values). The aim is to estimate these parameters by finding the set of parameters ***θ*** that minimizes the function *G*(***θ***), which describes the deviation between model output and experimental measurement data *y*:

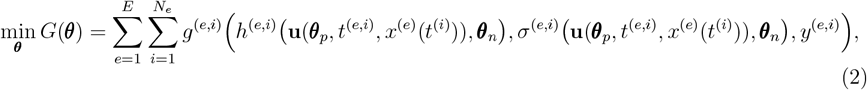

where the observable function *h* links the ODE system to the observed data, and *σ* describes the measurement noise formula. The problem is subject to upper and lower bounds **lb** ≤ ***θ*** ≤ **ub**. Note that the input and initial value functions can modify parameters that are part of the ODE system but not included in ***θ***. Additionally, the dynamic parameter can be condition-specific, specifically 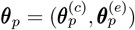, where *c* denotes constant between simulation/experimental conditions. In the PEtab standard, the function *g*^(*e,i*)^ and, in extension, *G* represent negative log-likelihoods. So given a normal measurement noise *g*^(*e,i*)^ would, dropping super-scripts and time *t* for ease of notation, look like;

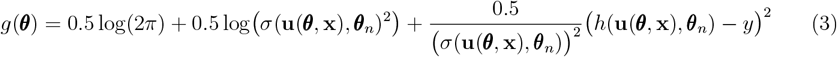

### Gradient computations

For small models, the gradient ∇*G* is traditionally computed using forward sensitivity analysis (with less than 100 parameters) and adjoint sensitivity analysis for larger models [30]. In addition, our PEtab importer can use forward-mode automatic differentiation to compute both the gradient and the Hessian [46, 1].

#### Computing ∇*G*(*θ*) via automatic differentiation

`PEtab.jl` computes the gradient via forward-mode automatic differentiation by propagating dual numbers, *x* + *yϵ* with *ϵ*^2^ = 0, through the program. When a differentiable function *g* : ℝ^*n*^ → ℝ is applied to a dual number with dual seed *y*, the dual part evaluates to the directional derivative ⟨∇*g*(*x*), *y*⟩. Thus, by running *n*-forward passes with the unit vectors as seeds, the gradient is obtained. It is further possible to employ forward mode AD for a vector function *g* : ℝ^*n*^ →ℝ^*m*^. Here, a single forward pass computes the Jacobian vector product **Jr** without building the Jacobian.

#### Implementation in this study

For computing gradients and Jacobians via forward-mode AD, `PEtab.jl` uses `ForwardDiff.jl` [46]. Note that `ForwardDiff.jl` uses multidimensional dual numbers, which can calculate multiple directional derivatives in a single forward pass. The number of derivatives computed per forward pass is referred to as the chunk size. Additionally, `ForwardDiff.jl` can compute the complete Hessian.

#### Computing ∇*G*(*θ*) via forward sensitivity equations

For ease of notation, as the gradient is a linear operator, we derive ∇_*p*_*G* for the following expression (dropping parameters not part of the ODE system and input function *x*):

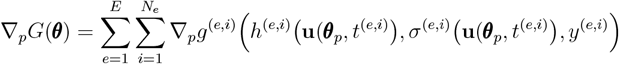

Via the chain rule

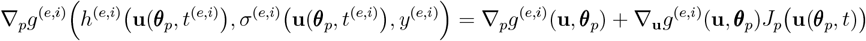

where 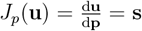 are the sensitivities, and via the chain-rule ∇_*p*_*g*^(*e,i*)^(**u, *θ***_*p*_) for parameter *j* equals (similar holds for derivative with respect to **u**)

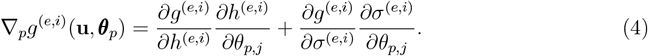

Both ∇_*p*_*g*^(*e,i*)^(**u**, *θ*_*p*_) and ∇_*u*_*g*^(*e,i*)^(**u, *θ***_*p*_) can be computed symbolically, while the sensitivities can be computed by solving the expanded ODE system;

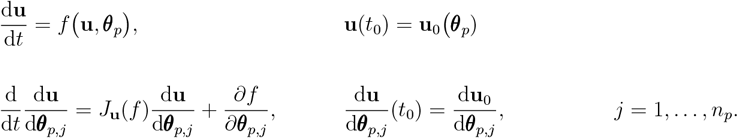

### Implementation in this study

`PEtab.jl` computes the sensitivities *J*_*p*_(**u**) via forward-mode AD (the Jacobian of the ODE solution). AMICI solves the expanded ODE system. The remaining derivatives ∇_*p*_*g* and ∇_**u**_*g* are computed symbolically.

#### Computing ∇*G*(*θ*) via adjoint sensitivity analysis

Here we derive an expression of the gradient for an experimental condition *e* (one forward simulation) denoted ∇*G*^(*e*)^, since the gradient is the sum of the gradients across experimental conditions 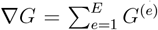 Further, for ease of notation we drop parameters not part of the ODE system. First, we introduce a Lagrangian multiplier with the same dimension as the state vector of the ODE, **u**; ***λ***(*t*) ∈ ℝ^*m*^, and rearrange the equation for *G*^(*e*)^ = *M* ^(*e*)^

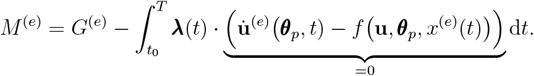

After the rearrangement, the gradient can be expressed as

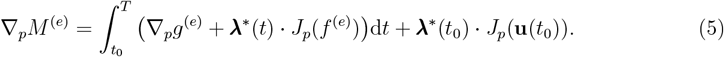

where *J*_*p*_(*f* ^(*e*)^) = *J*_*p*(_*f* (**u, *θ***_*p*_, *x*^*e*^(*t*))) is a Jacobian, and ***λ*** is the solution to

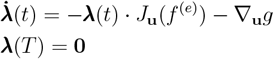

where *J*_**u**_(*f* ^(*e*)^) = *J*_**u**_ *f* (**u, *θ***_*p*_, *x*^*e*^(*t*)) is the Jacobian of the right-hand side of the ODE model. The sensitivities at time zero, 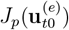, can be computed without solving the ODE system, and ∇_**u**_*g*^(*e*)^ and ∇_*p*_*g*^(*e*)^ have closed forms which can be computed symbolically (see above for forward sensitivities). The gradient can be computed by solving the ODE system for ***λ***, and then by evaluating the integral in Eq. (5). It should be noted that since we only observe data at discrete time-points, 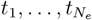 when solving for ***λ*** the term −∇_**u**_*g* is applied as a jump (discrete event) at the observed time-points. Further, following the PEtab standard *g* is time-point *i* dependent; *g*^(*e,i*)^.

### Implementation in this study

`PEtab.jl` can use either the quadrature and interpolation approach in `SciMLSensitivity.jl` [42, 34]. The vector-Jacobian-product ***λ***^*∗*^ *J*_*p*_(*f* ^(*e*)^) is computed in a Jacobian-free manner either via the Enzyme [39], or `ReverseDiff.jl` automatic differentiation libraries. AMICI employs the Sundial’s adjoint implementation [10], which is similar to the interpolation in `SciMLSensitivity.jl` [42]. Specifically, both approaches utilize an interpolation of **u**(*t*) to solve for ***λ***(*t*), and thus do not store the entire trajectory of ***λ***(*t*) over time. Both approaches employ a similar checkpointing strategy where the chosen interval points come from the forward solution, and when the reverse pass enters a new interval, the original ODE is re-solved on the interval [*t*_*k−*1_, *t*_*k*_]. It should also be noted that in AMICI, the Jacobian is computed symbolically, and the product is computed via sparse vector-matrix multiplication.

### Hessian computations

- BFGS. A Quasi-Newton method approximating the Hessian with a positive-definite matrix *B* updated iteratively from previous computation steps assumes *H*(*x*_*k*+1_) ≈ *H*(*x*_*k*_). Since the computation doesn’t require second-order sensitivities, the computational complexity is substantially reduced for large models. Let *f* be a twice differentiable (objective) function *f* : ℝ^*n*^ →ℝ. Note that the update scheme is specific to each Quasi-Newton method, but *B* needs to fulfil the Quasi-Newton condition in any case: *B*_*k*+1_[*x*_*k*+1_ −*x*_*k*_] = ∇ *f* (*x*_*k*+1_) − *f* (*x*_*k*_). The update for the Broyden-Fletcher-Goldfarb-Shannon (BFGS) algorithm is given by

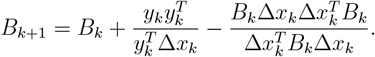
- Gauss-Newton. Similarly, the Gauss-Newton method avoids the calculation of second derivatives as well by recasting the Hessian computation as a non-linear least squares problem: 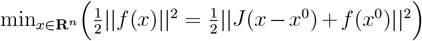. By linearization of the objective function, *f* through Taylor series expansion of order one and using the Jacobian matrix *J*, the Hessian (after some calculus) can be approximated by *J*(*x*) + *H*(*x*)Δ*x* = 0 avoiding computing and inverting a potentially large Hessian matrix as a limiting factor. The Gauss-Newton matrix approximating the Hessian *H* is given by:

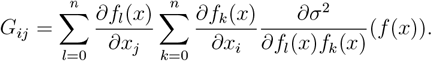

### Robustness of results

All Julia benchmarks were performed using Julia version 1.8.5, except the stochastic ones, which (due to being carried out at a later stage) were performed on version 1.10.2 [3]. The `pyPESTO` and AMICI benchmarks were performed using Python version 3.10.6, Amici version 0.17 to address bugs for Smith and Schwen models [10] and version 0.15 for remaining benchmarks, and `pyPESTO` version 0.2.15 [49]. The code used to perform all benchmarks can be found at https://github.com/cvijoviclab/PEtab_benchmark. This repository also provides a comprehensive guide on how to set up the necessary environment for running the benchmarks. It should be noted that for the benchmarks, we used a predecessor version of `PEtab.jl`, which was later migrated to a new repository to create the formal Julia package repository. This holds for all models, except Bruno, where due to a bug in the `PEtab.jl` predecessor version found during revision, we ran the benchmark with `PEtab.jl` version 2.14. Complementary, identical benchmarks have been run on a second compute infrastructure to demonstrate the robustness and reproducibility of results corresponding to the Julia benchmarks with Julia version 1.9.0 respectively, `pyPESTO` and AMICI benchmarks with Python 3.10.4, AMICI version 0.15 and `pyPESTO` 0.3.0.

### Disclosure - usage of AI-assisted tools

We used ChatGTP4 solely for text editing to improve the readability of the manuscript. Specifically, once the first draft, including references, had been completed, for each paragraph, we ran the ChatGTP4 with the prompt: *Please improve the readability of this paragraph*. Subsequently, the text produced by ChatGTP4 was edited to avoid any potential inaccuracies and misinterpretations or to avoid unclear formulations. To further minimise any chance for potential inaccuracies from the language model, ChatGTP4 was applied prior to circulating the manuscript between all the authors. Taken together, the authors reviewed and edited the content as needed and took full responsibility for the content of the publication.

## Acknowledgements

This work was supported by the Swedish Research Council (VR2017-05117 and VR2023-04319) and the Swedish Foundation for Strategic Research (FFL15-0238) to MC. FF was supported by the Francis Crick Institute, which receives its core funding from Cancer Research UK (CC2242), the UK Medical Research Council (CC2242), and the Wellcome Trust (CC2242). JH and SG are supported by the German Research Foundation (CRC 1454 - Metaflammation, project no. 432325352 and AMICI, project no. 443187771). JH acknowledges funding by the German Research Foundation under Germany’s Excellence Strategy EXC 2151 - 390873048 (ImmunoSensation2) and EXC 2047 - 390685813 (Hausdorff Center for Mathematics) and financial support via a Schlegel Professorship at the University of Bonn. This material is based upon work supported by the National Science Foundation under grant no. DMS-2325184 to TL. For the purpose of Open Access, the author F.F. has applied a CC BY public copyright licence to any Author Accepted Manuscript version arising from this submission.

## Author contributions

S.P. and M.C. conceptualization; S.P., D.O., V.H. software; S.P., F.F, S.G, D.O., V.H., T.L formal analysis and validation; S.P. visualization; All authors wrote and edited the manuscript. J.H. and M.C. supervised the research.

