## Supplementary Text for "PEtab.jl: Advancing the Efficiency and Utility of Dynamic Modelling"

#### Extended discussion on ODE solvers

In the benchmark evaluating a selection of ODE solvers on smaller models with randomly chosen parameters (Fig. 2 and S1), for a subset of models and parameters, the ODE solvers encountered simulation failure. To investigate why, we investigated the ODE solver return codes (Fig. S2a). The non-stiff, explicit ODE solvers (Vern7 and Tsit5) primarily failed with code *Maxiters*, which is common when using a non-stiff solver on a stiff problem.

Regarding the multi-step BDF solvers QNDF and CVODE_BDF, the former failed more frequently (Fig. 2). The most common error code for QNDF was *DtLessThanMin* which means it fails to meet integration tolerances, and as this happened for times close to zero the results suggests it fails during the startup phase where it relies on lower-order methods. For a subset of SIR models (Bertozzi and Giordano) both solvers had code *Maxiters*, which can be fixed by reducing solver tolerances (Fig. S2b), and probably by setting larger maximum iteration number. Overall, given that CVODE_BDF and QNDF have comparable runtimes but CVODE_BDF is more reliable (Fig. 2), and there are models (e.g. Schwen) where lowering tolerances for QNDF do not make it able to simulate for all vectors that CVODE_BDF can, for simulation-based tasks with BDF solvers we recommend CVODE_BDF. Nevertheless, in cases where QNDF performs well (Bachmann Fig. 2 and Fig. 3), as it is compatible with efficient automatic differentiation (Fig. 4), for gradient-based tasks, QNDF can be a good choice.

The composite solver Vern7Rodas5P failed more frequently for stiffer models (Schwen, Bachmann, Weber, and Fiedler) than CVODE_BDF, often with error code *Unstable*. This suggests the composite solver does not switch to the stiff solver in time for these models, and in addition, for these models (except Weber), lowering the tolerance did not make it able to simulate the model. Conversely, for models with less apparent stiffness, like the SIR models (Bertozzi, Zhao, Okunghae, and Giordano), Vern7Rodas5P was the fastest solver.

The Rosenbrock solver Rodas5P was reliable, and it only had more simulation failures than CVODE_BDF for two models (Borghans 80 vs 77, Fiedler 8 vs 0). It had the *unstable* error code for both the Fiedler and Giordano models, but for the latter, it could simulate the model with lower tolerances (Fig. S2a). Overall, as Rodas5P generally is more accurate than BDF methods (Fig. S1a) and often is faster for models with less than 25 ODEs (Fig. 2), we recommend Rosenbrock solvers like Rodas5P for smaller models. Moreover, the Fiedler model was the only case where Rodas5P had problems compared to other solvers, but the results suggest that in cases where Rosenbrock solvers often have recode *Unstable*, switching to a pdf-solver can help.

Lastly, Borghans (a spiking model) and Crauste (a virus growth model) had many simulation failures. Error codes indicate unstable dynamics for a subset of parameters, and reducing tolerances often failed to resolve the issue. Importantly, the composite solver had a few error-codes of *Longtime* (solver was aborted as it took > 15min), potentially because it diverged so badly it ended in a part of model phase-space where solving linear systems takes considerable time. Regardless, this suggests that for numerically challenging models avoid composite solvers as they can put workflows to a halt with long simulation times, and even though stiff solvers performed better here, there is no clear best choice.

In a flow-chart (Fig. 6), we summarise the result and make the following recommendations: for less stiff models, such as many SIR models, composite solvers, for smaller, stiffer models, like mechanistic models, Rosenbrock solvers like Rodas5P. Should Rosenbrock solvers have problems (e.g. Fiedler model), switching to a BDF method can help. For larger, stiffer models, BDF solvers are suitable. If issues occur with QNDF, CVODE_BDF is reliable. For certain large models like those with many events, SDIRK solvers like KenCarp4 should also be considered (Fig. 3).

#### Extended discussion on parameter estimation results

In the parameter estimation benchmark results for the Fides optimiser with the Gauss-Newton Hessian approximation for a subset of models (e.g. Crauste and Isensee) `pyPESTO` and `PEtab.jl` have a noticeable difference in the number of converged runs (Fig. 7). As both pyPESTO and PEtab.jl are thoroughly tested (further confirmed by comparing derivatives between software), the variance in converged runs is likely because of it it being hard to accurately compute the Gauss-Newton Hessian approximation for a subset of numerically challenging models. This is further supported by looking closer at the Isensee and Crauste models where number of converged runs differ, compared to the Boehm model where both software have similar number of converged runs (Tab. S1).

The Isensee model is large, stiff, and has a pre-equilibration condition (steady-state simulations). Just like for the other pre-equilibration models (Zheng, Weber and Brannmark), neither software had many converged runs. Moreover, for the Isensee model AMICI's sophisticated approach for steady-state handling, which for example was used used for the Zheng model, cannot be employed as the model has a singular Jacobian. Overall, the stiffness, size and pre-equilibration makes it challenging to compute derivatives. For example, the difference in $\ell_2$-norm between a high accuracy gradient and Gauss-Newton Hessian (obtained low solver tolerances), is larger than for the Boehm model (Tab. S1). Further, the differences are larger for the Gauss-Newton Hessian than the gradient (Tab. S1). Naturally, one action to obtain more accurate gradients in practice could be to lower ODE solver tolerances, however, this also increases the number of simulation failures [?].

The Crauste model is even more numerically challenging than the Isensee model. Lowering ODE solver tolerances results in drastic differences for both the gradient and Guass-Newton Hessian approximation compared to the high accuracy gradient reference. The difference is larger for the Gauss-Newton Hessian (Tab. S1).

In summary, the variance in converged when running parameter estimation using Fides with the Gauss-Newton Hessian approximation for a subset of models (e.g. Crauste and Isensee), is likely because computing gradients, and particular a Hessian approximation is challenging for a subset of models. Hence, using different gradient methods and ODE solvers likely affects the results. Given that neither `PEtab.jl` nor `pyPESTO` consistently shows higher convergence rates across models, neither software with respect to this aspect cannot really be recommended over the other, and identifying exactly why for example `PEtab.jl` performed better for Iseense is an interesting future research direction. Lastly, the difference in converged runs is smaller when running Fides with BFGS Hessian approximation, likely because BFGS uses gradients across the history for a robust estimate Hessian estimate, and it only uses the gradient from PEtab.jl and pyPESTO.

### Supplementary tables

**Table 1: Effect of changing ODE solver tolerance on gradient and Gauss-Newton Hessian (H) accuracy**. For Crauste and Boehm the reference gradient and Hessian were computed using tolerance = $10^{-12}$, while due to simulation failure the Isensee the tolerance $10^{-11}$ was used. The $\ell_2$ columns are the difference in $\ell_2$ norm compared to the reference, and the max-diff columns are the maximum difference in an element compared to the reference.

| Model | Solver tolerance | $\ell_2$-norm grad. | $\ell_2$-norm H. | Max. grad. diff | Max. H diff |
|---|---|---|---|---|---|
| Crauste | $10^{-6}$ | $5.9 \times 10^4$ | $4.8 \times 10^{11}$ | $4.2 \times 10^4$ | $2.5 \times 10^{11}$ |
| Crauste | $10^{-8}$ | 492 | $3.9 \times 10^9$ | 350 | $2 \times 10^9$ |
| Crauste | $10^{-10}$ | 5.6 | $4.6 \times 10^7$ | 4 | $2.3 \times 10^7$ |
| Boehm | $10^{-6}$ | $4.6 \times 10^{-5}$ | $1.9 \times 10^{-4}$ | $3.7 \times 10^{-5}$ | $7.9 \times 10^{-5}$ |
| Boehm | $10^{-8}$ | $1.1 \times 10^{-6}$ | $4.4 \times 10^{-6}$ | $8.7 \times 10^{-7}$ | $1.8 \times 10^{-8}$ |
| Boehm | $10^{-10}$ | $1.7 \times 10^{-8}$ | $1.4 \times 10^{-8}$ | $1.7 \times 10^{-8}$ | $2.2 \times 10^{-8}$ |
| Isensee | $10^{-6}$ | 1.5 | 426 | 1.18 | 173 |
| Isensee | $10^{-8}$ | 0.008 | 2.0 | 0.006 | 0.81 |
| Isensee | $10^{-10}$ | $7.5 \times 10^{-5}$ | 0.01 | $4.4 \times 10^{-5}$ | 0.004 |

**Table 2: SBML component and test tags supported by SBMLImporter.jl.** For comparison, AMICI is also included. Tags apply to the semantic test suite. Only tags where either tool passes one test with the tag are reported in the table. To date, AMICI passes 1247 and SBMLImporter.jl 1257 of the 1821 test cases. The main differences boil down to AMICI supporting hierarchical models, while SBMLImporter.jl supports StoichiometryMath.

| Tag | Tag type | Supported AMICI | Supported SBMLImporter.jl |
|---|---|---|---|
| AlgebraicRule | Component tag | × | × |
| AssignmentRule | Component tag | × | × |
| comp | Component tag | × | |
| Compartment | Component tag | × | × |
| CSymbolAvogadro | Component tag | × | × |
| CSymbolRateOf | Component tag | × | × |
| CSymbolTime | Component tag | × | × |
| Deletion | Component tag | × | × |
| EventNoDelay | Component tag | × | × |
| ExternalModelDefinition | Component tag | × | |
| FunctionDefinition | Component tag | × | × |
| InitialAssignment | Component tag | × | × |
| ModelDefinition | Component tag | × | |
| Parameter | Component tag | × | × |
| Port | Component tag | × | |
| RateRule | Component tag | × | × |
| Reaction | Component tag | × | × |
| ReplacedBy | Component tag | × | |
| ReplacedElement | Component tag | × | |
| SBaseRef | Component tag | × | |
| Species | Component tag | × | × |
| StoichiometryMath | Component tag | | × |
| Submodel | Component tag | × | |
| 0D-Compartment | Test tag | × | × |
| Amount | Test tag | × | × |
| AssignedConstantStoichiometry | Test tag | × | × |
| AssignedVariableStoichiometry | Test tag | × | × |
| BoolNumericSwap | Test tag | × | × |
| BoundaryCondition | Test tag | × | × |
| comp | Test tag | × | |
| Concentration | Test tag | × | × |
| ConstantSpecies | Test tag | × | × |
| ConversionFactors | Test tag | × | × |
| DefaultValue | Test tag | × | × |
| EventT0Firing | Test tag | × | × |
| ExtentConversionFactor | Test tag | × | |
| HasOnlySubstanceUnits | Test tag | × | × |
| InitialValueReassigned | Test tag | × | × |
| L3v2MathML | Test tag | × | × |
| LocalParameters | Test tag | × | × |
| MultiCompartment | Test tag | × | × |
| NoMathML | Test tag | × | × |
| NonConstantCompartment | Test tag | × | × |

| | | | |
|---|---|---|---|
| NonConstantParameter | Test tag | × | × |
| NonUnityCompartment | Test tag | × | × |
| NonUnityStoichiometry | Test tag | × | × |
| ReversibleReaction | Test tag | × | × |
| SpeciesReferenceInMath | Test tag | × | × |
| SubmodelOutput | Test tag | × | |
| TimeConversionFactor | Test tag | × | |
| UncommonMathML | Test tag | × | × |
| VolumeConcentrationRates | Test tag | × | × |

**Table 3: Runtime evaluation for the cost (objective) and gradient**. Benchmark is performed at reported values. AMICI uses CVODE-BDF, while the Julia PEtab importer uses the Rodas5P and QNDF ODE solvers. Ratio represents the ratio between the runtime for the gradient and cost, and Inf denotes gradient computational failure. The number of model parameters the gradient is taken to is given in the rightmost column.

| Solver | Time cost [s] | Time gradient [s] | Ratio | Number of parameters |
|---|---|---|---|---|
| Zheng_PNAS2012 | | | | |
| QNDF | 7.05e-04 | 3.82e-02 | 5.41e+01 | 46 |
| Rodas5P | 6.90e-04 | 4.93e-02 | 7.14e+01 | 46 |
| AMICI | 2.84e-04 | 1.43e-02 | 5.03e+01 | 46 |
| Weber_BMC2015 | | | | |
| QNDF | 1.48e-03 | 2.04e-02 | 1.38e+01 | 36 |
| Rodas5P | 2.55e-03 | 2.97e-02 | 1.16e+01 | 36 |
| AMICI | 1.15e-03 | 6.19e-02 | 5.36e+01 | 36 |
| Sneyd_PNAS2002 | | | | |
| QNDF | 2.84e-03 | 1.83e-02 | 6.47e+00 | 15 |
| Rodas5P | 2.38e-03 | 1.85e-02 | 7.77e+00 | 15 |
| AMICI | 2.47e-03 | 4.82e-02 | 1.95e+01 | 15 |
| Schwen_PONE2014 | | | | |
| QNDF | 7.98e-03 | 6.11e-02 | 7.65e+00 | 30 |
| Rodas5P | 8.81e-03 | 7.25e-02 | 8.23e+00 | 30 |
| AMICI | 5.72e-03 | 1.43e-01 | 2.51e+01 | 30 |
| Rahman_MBS2016 | | | | |
| QNDF | 2.75e-04 | 1.02e-03 | 3.71e+00 | 9 |
| Rodas5P | 4.62e-04 | 1.48e-03 | 3.21e+00 | 9 |
| AMICI | 2.95e-04 | 5.33e-03 | 1.81e+01 | 9 |
| Oliveira_NatCommun2021 | | | | |
| QNDF | 1.06e-03 | Inf | Inf | 12 |
| Rodas5P | 5.65e-04 | 4.42e-03 | 7.82e+00 | 12 |
| AMICI | 7.53e-04 | 8.25e-03 | 1.10e+01 | 12 |
| Okuonghae_ChaosSolitonsFractals2020 | | | | |
| QNDF | 3.22e-04 | 1.64e-03 | 5.10e+00 | 16 |
| Rodas5P | 2.11e-04 | 1.52e-03 | 7.23e+00 | 16 |

| | | | | |
|---|---|---|---|---|
| AMICI | 3.13e-04 | 5.83e-03 | 1.86e+01 | 16 |
| Lucarelli_CellSystems2018 | | | | |
| QNDF | 1.44e-02 | 2.28e+00 | 1.59e+02 | 84 |
| Rodas5P | 4.58e-02 | 1.04e+01 | 2.26e+02 | 84 |
| AMICI | 1.02e-02 | 1.16e+00 | 1.13e+02 | 84 |
| Isensee_JCB2018 | | | | |
| QNDF | 1.02e-01 | 2.94e+00 | 2.87e+01 | 46 |
| Rodas5P | 1.07e-01 | 3.74e+00 | 3.51e+01 | 46 |
| AMICI | 6.09e-02 | 2.99e+00 | 4.91e+01 | 46 |
| Fujita_SciSignal2010 | | | | |
| Rodas5P | 3.06e-03 | 2.32e-02 | 7.59e+00 | 19 |
| AMICI | 3.26e-03 | 1.28e-01 | 3.92e+01 | 19 |
| Fiedler_BMC2016 | | | | |
| QNDF | 7.36e-04 | 4.40e-03 | 5.98e+00 | 22 |
| Rodas5P | 6.77e-04 | 6.07e-03 | 8.97e+00 | 22 |
| AMICI | 7.79e-04 | 2.29e-02 | 2.94e+01 | 22 |
| Elowitz_Nature2000 | | | | |
| QNDF | 2.59e-03 | 1.91e-02 | 7.38e+00 | 21 |
| Rodas5P | 3.66e-03 | 2.73e-02 | 7.46e+00 | 21 |
| AMICI | 1.61e-03 | 4.04e-02 | 2.50e+01 | 21 |
| Crauste_CellSystems2017 | | | | |
| QNDF | 8.97e-04 | 5.26e-03 | 5.86e+00 | 12 |
| Rodas5P | 8.53e-04 | 6.87e-03 | 8.06e+00 | 12 |
| AMICI | 6.48e-04 | 1.40e-02 | 2.16e+01 | 12 |
| Bruno_JExpBot2016 | | | | |
| QNDF | 5.02e-04 | 3.22e-03 | 6.41e+00 | 13 |
| Rodas5P | 3.56e-04 | 2.32e-03 | 6.50e+00 | 13 |
| AMICI | 7.13e-04 | 6.89e-03 | 9.67e+00 | 13 |
| Brannmark_JBC2010 | | | | |
| QNDF | 2.75e-03 | 2.23e-02 | 8.11e+00 | 22 |
| Rodas5P | 2.98e-03 | 2.48e-02 | 8.33e+00 | 22 |
| AMICI | 3.35e-03 | 8.14e-02 | 2.43e+01 | 22 |
| Borghans_BiophysChem1997 | | | | |
| QNDF | 6.94e-03 | 5.88e-02 | 8.47e+00 | 23 |
| Rodas5P | 1.32e-02 | 1.23e-01 | 9.29e+00 | 23 |
| AMICI | 2.31e-03 | 7.11e-02 | 3.08e+01 | 23 |
| Boehm_JProteomeRes2014 | | | | |
| QNDF | 5.69e-04 | 1.74e-03 | 3.06e+00 | 9 |
| Rodas5P | 5.88e-04 | 1.94e-03 | 3.30e+00 | 9 |
| AMICI | 4.46e-04 | 4.45e-03 | 9.98e+00 | 9 |
| Beer_MolBioSystems2014 | | | | |

| | | | | |
|---|---|---|---|---|
| QNDF | 1.07e-02 | 8.95e-04 | 8.33e-02 | 72 |
| Rodas5P | 1.09e-02 | 3.04e-01 | 2.78e+01 | 72 |
| AMICI | 2.22e-02 | 1.79e-01 | 8.05e+00 | 72 |
| Bachmann_MSB2011 | | | | |
| QNDF | 3.22e-02 | 7.46e-01 | 2.32e+01 | 113 |
| Rodas5P | 5.48e-02 | 1.54e+00 | 2.81e+01 | 113 |
| AMICI | 1.83e-02 | 1.08e+00 | 5.92e+01 | 113 |

**Table 4: Runtime evaluation for the cost (objective) and Hessian**. Benchmark is performed at reported values. Julia PEtab importer uses the Rodas5P and QNDF ODE solvers. The ratio represents the ratio between the runtime for the Hessian and cost, and Inf denotes Hessian computation failure. The number of model parameters the Hessian is taken to is given in the rightmost column. Note the Hessian is computed with a forward-over-forward approach and thus should theoretically have a quadratic scaling with the number of parameters.

| Solver | Time Hessian [s] | Time cost [s] | Ratio | Number of parameters |
|---|---|---|---|---|
| Zheng_PNAS2012 | | | | |
| QNDF | 3.63e+00 | 8.92e-04 | 4.07e+03 | 46 |
| Rodas5P | 5.90e+00 | 8.60e-04 | 6.87e+03 | 46 |
| Weber_BMC2015 | | | | |
| QNDF | 1.98e+00 | 1.82e-03 | 1.09e+03 | 36 |
| Rodas5P | 5.89e+00 | 2.85e-03 | 2.07e+03 | 36 |
| Sneyd_PNAS2002 | | | | |
| QNDF | 9.48e-01 | 3.48e-03 | 2.73e+02 | 15 |
| Rodas5P | 1.45e+00 | 2.54e-03 | 5.69e+02 | 15 |
| Schwen_PONE2014 | | | | |
| QNDF | 3.70e+00 | 7.93e-03 | 4.66e+02 | 30 |
| Rodas5P | 5.55e+00 | 8.64e-03 | 6.42e+02 | 30 |
| Rahman_MBS2016 | | | | |
| QNDF | 1.42e-02 | 2.71e-04 | 5.22e+01 | 9 |
| Rodas5P | 4.85e-02 | 3.38e-04 | 1.44e+02 | 9 |
| Oliveira_NatCommun2021 | | | | |
| QNDF | 1.40e-03 | Inf | Inf | 12 |
| Rodas5P | 7.42e-02 | 5.68e-04 | 1.31e+02 | 12 |
| Okuonghae_ChaosSolitonsFractals2020 | | | | |
| QNDF | 2.78e-02 | 1.96e-04 | 1.42e+02 | 16 |
| Rodas5P | 3.89e-02 | 1.64e-04 | 2.37e+02 | 16 |
| Lucarelli_CellSystems2018 | | | | |
| QNDF | 3.30e+02 | 2.04e-02 | 1.62e+04 | 84 |
| Rodas5P | 1.50e+03 | 6.08e-02 | 2.47e+04 | 84 |
| Isensee_JCB2018 | | | | |
| Rodas5P | Inf | 1.37e-01 | Inf | 46 |
| Fujita_SciSignal2010 | | | | |
| QNDF | 9.62e-01 | 4.58e-03 | 2.10e+02 | 19 |
| Rodas5P | 1.05e+00 | 3.37e-03 | 3.13e+02 | 19 |
| Fiedler_BMC2016 | | | | |
| QNDF | 2.35e-01 | 9.11e-04 | 2.58e+02 | 22 |
| Rodas5P | 4.11e-01 | 8.86e-04 | 4.64e+02 | 22 |

| | | | | |
|---|---|---|---|---|
| Elowitz_Nature2000 | | | | |
| QNDF | 5.92e-01 | 2.72e-03 | 2.18e+02 | 21 |
| Rodas5P | 1.00e+00 | 3.91e-03 | 2.57e+02 | 21 |
| Crauste_CellSystems2017 | | | | |
| QNDF | 1.11e-01 | 1.14e-03 | 9.79e+01 | 12 |
| Rodas5P | 1.84e-01 | 9.79e-04 | 1.88e+02 | 12 |
| Bruno_JExpBot2016 | | | | |
| QNDF | 5.69e-02 | 6.14e-04 | 9.26e+01 | 13 |
| Rodas5P | 5.34e-02 | 3.77e-04 | 1.41e+02 | 13 |
| Brannmark_JBC2010 | | | | |
| QNDF | 1.08e+00 | 3.33e-03 | 3.25e+02 | 22 |
| Rodas5P | 1.88e+00 | 3.23e-03 | 5.80e+02 | 22 |
| Borghans_BiophysChem1997 | | | | |
| QNDF | 1.92e+00 | 6.79e-03 | 2.82e+02 | 23 |
| Rodas5P | 5.42e+00 | 1.30e-02 | 4.19e+02 | 23 |
| Boehm_JProteomeRes2014 | | | | |
| QNDF | 2.45e-02 | 6.69e-04 | 3.66e+01 | 9 |
| Rodas5P | 4.09e-02 | 6.93e-04 | 5.90e+01 | 9 |
| Beer_MolBioSystems2014 | | | | |
| Rodas5P | 3.31e+01 | 1.45e-02 | 2.29e+03 | 72 |
| Bachmann_MSB2011 | | | | |
| QNDF | 9.46e+02 | 3.86e-02 | 2.45e+04 | 113 |
| Rodas5P | 2.44e+03 | 7.05e-02 | 3.46e+04 | 113 |

### Supplementary figures

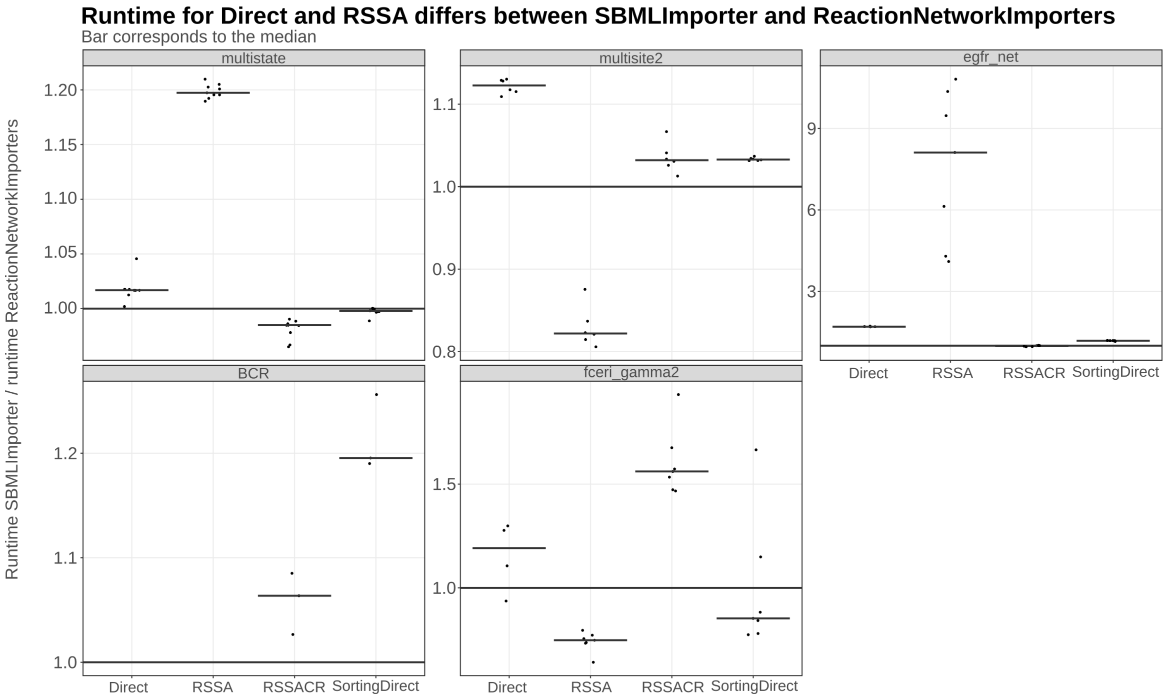


**Figure 1: Benchmark of stochastic simulators**: Simulation runtime for models imported as SBML files using `SBMLImporter.jl` divided by simulation runtime when the model is imported as .net files via `ReactionNetworkImporters.jl`, for the same simulation intervals as in Fig. 2. The order of reactions differs in the parsed Catalyst reaction network differs between the importers, which notably impacted runtime for Direct and RSSA methods; for instance, RSSA was about 1.2 times faster when using .net file imports for the multistate model. Before benchmarking, it was verified that the reaction and model parameters were identical between the importers.

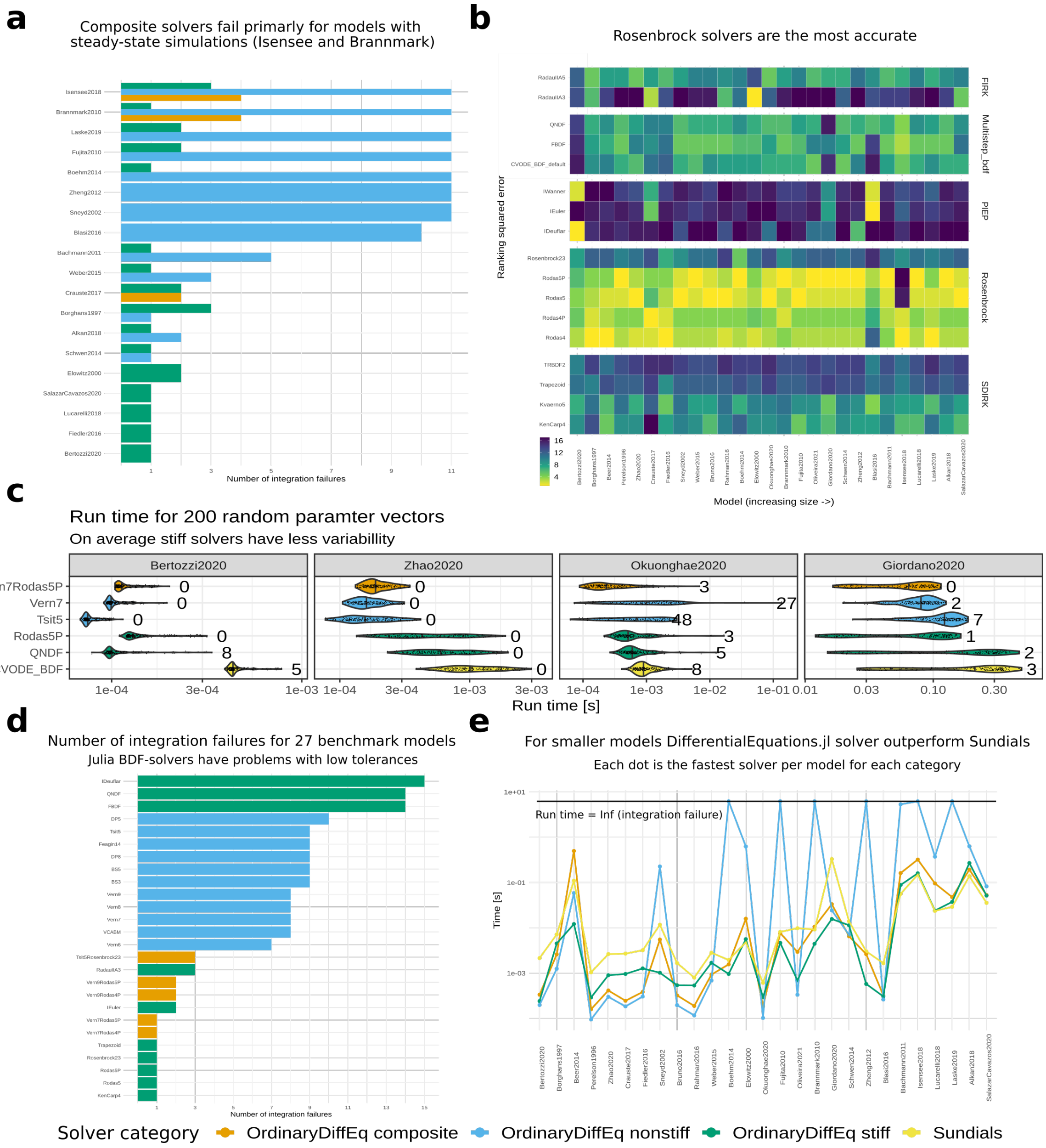


**Figure 2: Benchmark of Sundial and DifferentialEquations ODE solvers for 27 models. a)** Integration failures per model for a large selection of stiff solvers when using $abstol = reltol = 1 \times 10^{-8}$ **b)** Accuracy ranking per model for a large selection of stiff solvers grouped into their respective solver family when using the tolerances in a). A bright yellow color denotes a good ranking. Accuracy was computed by taking the $L_2$-norm for all time points with observed data for the relevant solver against a high accuracy solution computed with the Rodas5P with tolerances $abstol = reltol = 1 \times 10^{-15}$. **b)** Run-time across 100 random parameter vectors for 4 SIR models using non-stiff, stiff, and composite solvers that performed well. The centre line of each violin plot denotes the median, and the number at the end of the violin shows the number of integration failures for that solver. **d)** Total number of integration failures across all models with the tolerances when using the tolerances $abstol = 1 \times 10^{-16}$ and $reltol = 1 \times 10^{-8}$ b). **e)** Best average run-time (5 repetitions) for each solver category for all benchmark models using the tolerances in d).

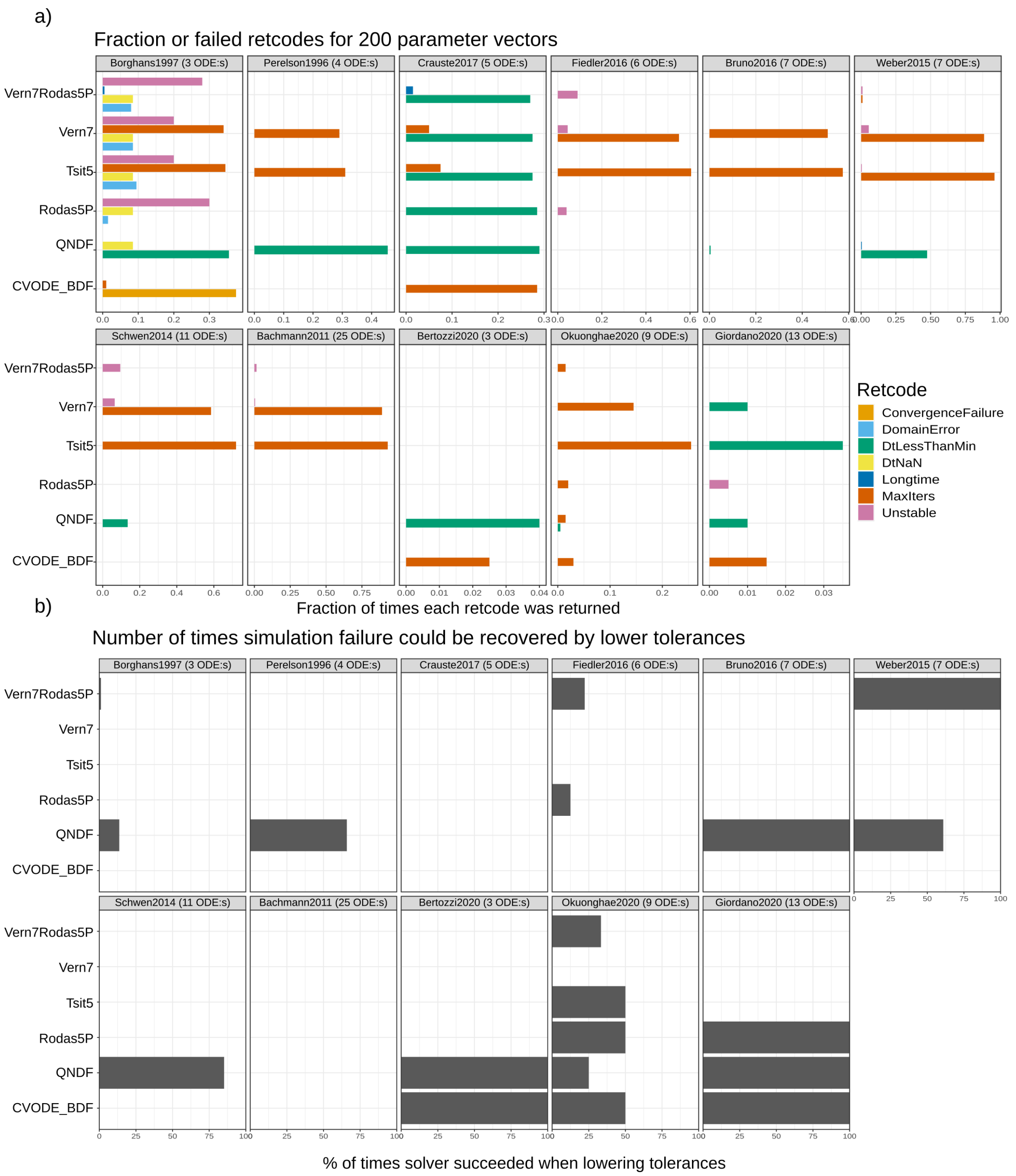


**Figure 3: ODE solver error codes when testing solvers for random parameters**. **a)** Fraction number of times each error appeared when simulating $n = 200$ random parameter vectors in Fig. 2 and Fig. S1. For example, for the Borghans model, CVODE_BDF failed for around $0.35 * 200 = 70$ cases with error code ConvergenceFailure. Only models where simulation failures occurred are included, so the Zhao model is excluded from the figure. **b)** Percentage of times simulation could be recovered by reducing ODE solver tolerances from $abstol = reltol = 1 \times 10^{-8}$ in steps of 10 to $abstol = reltol = 1 \times 10^{-4}$. For example, for the Bertozzi model, all simulation failures for CVODE_BDF could be resolved by increasing solver tolerances.

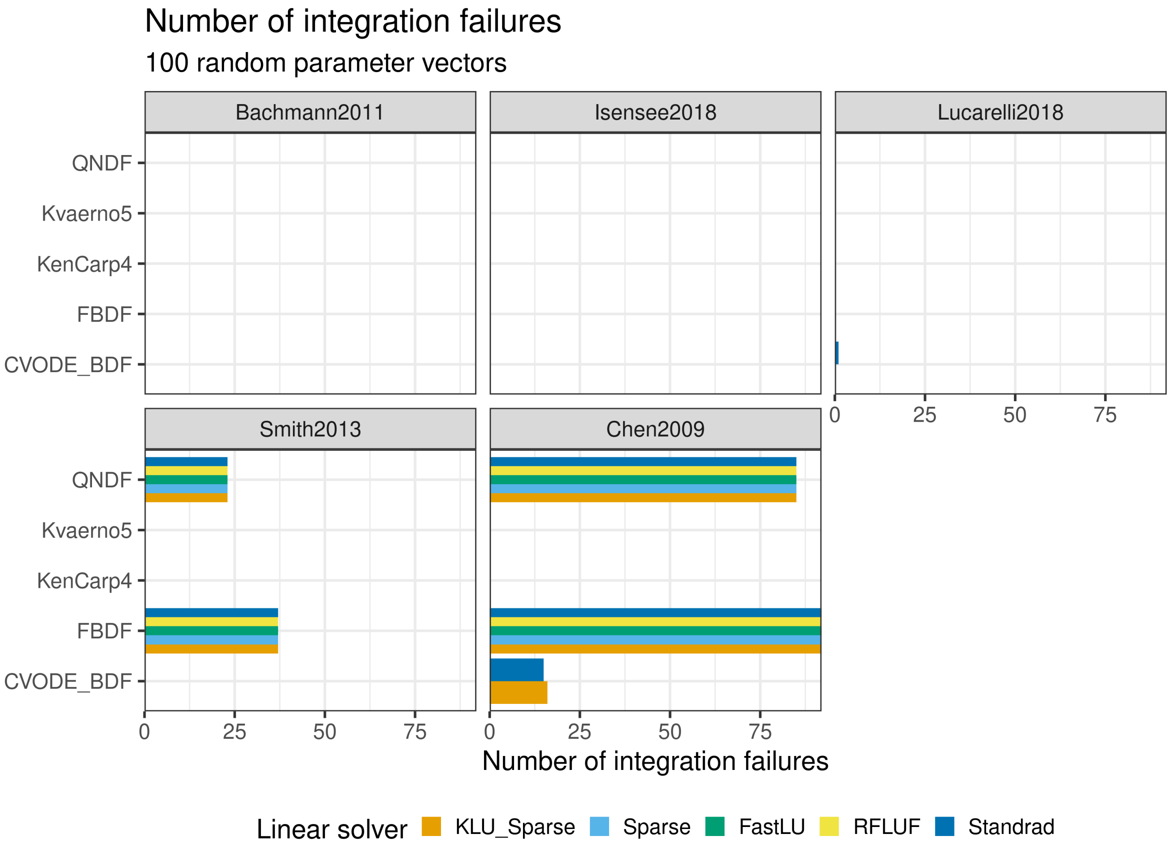


**Figure 4: Integration failures when trying ODE solvers for large models.** Number of integration failures for the 100 random parameter vectors in Fig. 3. Noticeably, the DifferentialEquations BDF solvers (QNDF and FBDF) frequently fail for the two bigger Smith and Chen models.

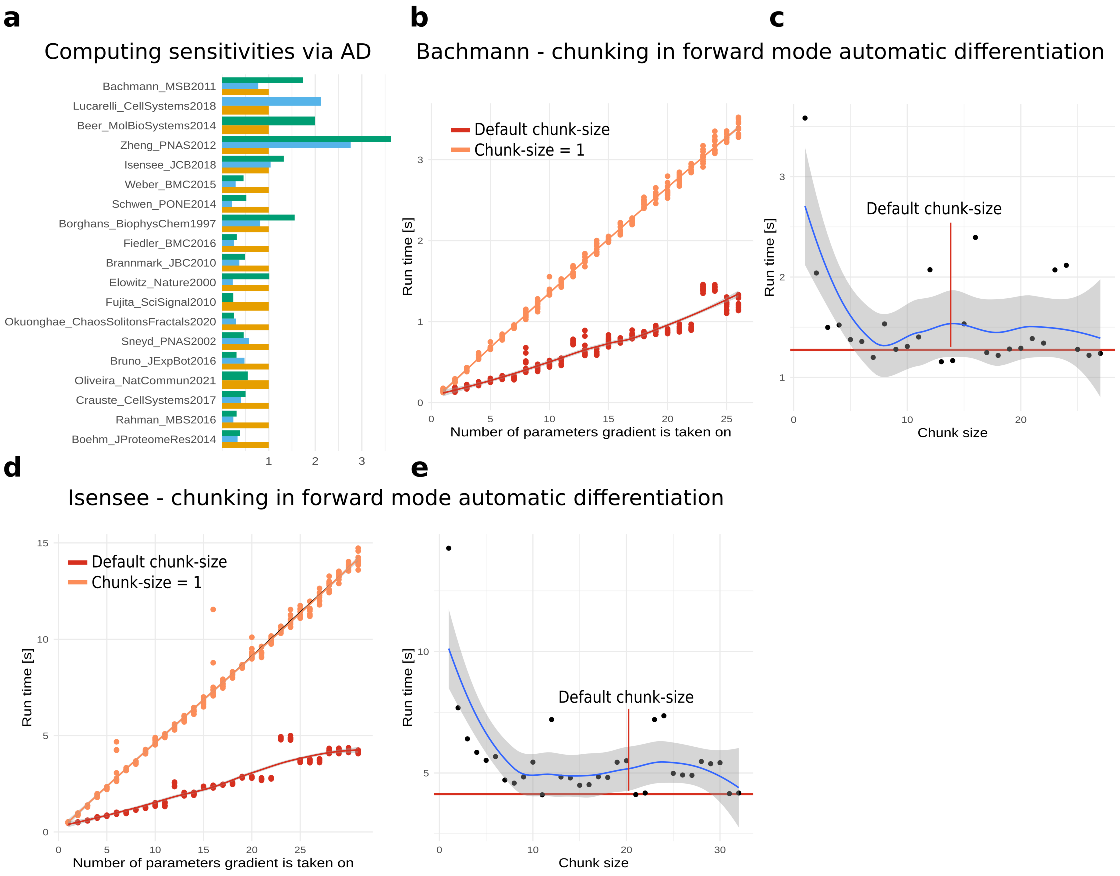


**Figure 5: Gradient benchmark for smaller models.** **a)** Run-time evaluation for the gradient at reported parameter values. AMICI computes the gradient by solving for the sensitivities via an expanded ODE system, while the Julia importer solves for the sensitives via Forward-mode AD. Run times are normalized with respect to AMICI. **b)** Gradient run-time for the Bachmann model when increasing the number of parameters using forward-mode AD without chunking and with the default number of chunks. **c)** Gradient run-time for the Bachmann model when taking the gradient on all parameters and testing all chunk sizes. **d-e** As in b-c) but for the Isensee model

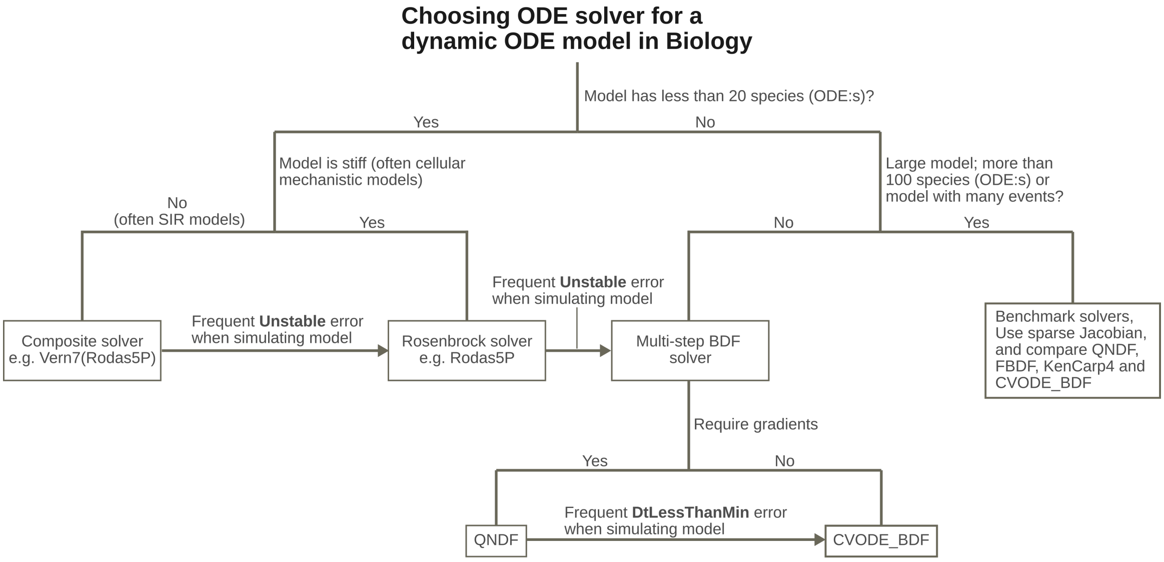


**Figure 6: Flowchart for ODE solver choice for dynamic models in biology**. This is a general recommendation based on the benchmarks (Fig. 2, 3, S1, S2), but it should be kept in mind that every problem is unique, and even though the settings here generally work well, they might not be optimal. For larger models (right part), we recommend benchmarking different solvers, as the runtime is substantial in this regime, so getting the best performance is extra important.

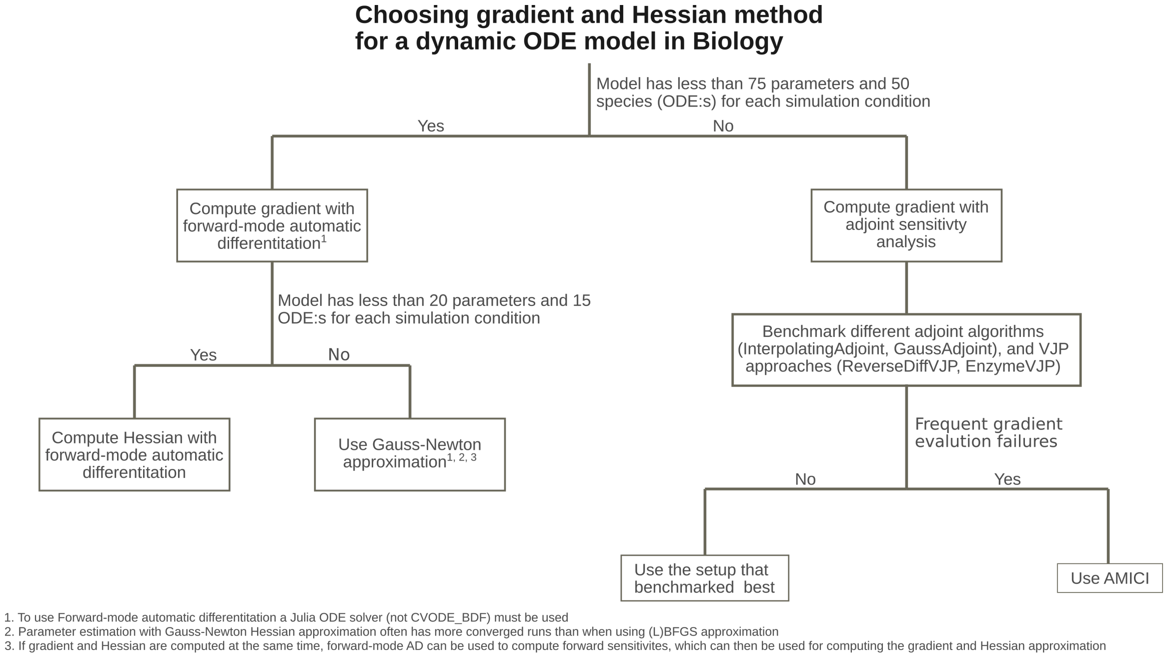


**Figure 7: Flowchart for gradient and Hessian method choice for dynamic models in biology**. This is a general recommendation based on the benchmarks (Fig. 3, 4), but it should be kept in mind that any problem is unique, and even though the settings here generally work well, they might not be optimal and/or change in the future. For larger models (right part), we recommend benchmarking different methods, as the runtime is substantial in this regime, getting the best performance is extra important.

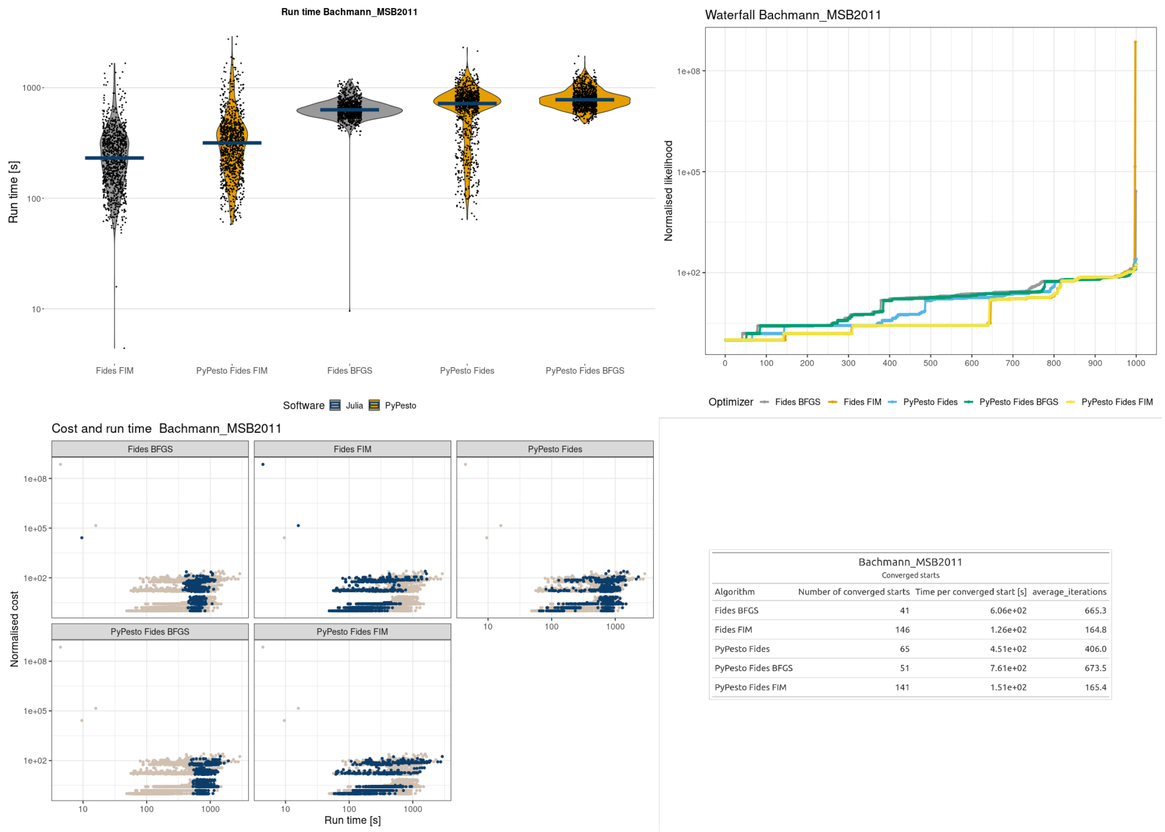


Bachmann_MSB2011

Converged starts

| Algorithm | Number of converged starts | Time per converged start [s] | average_iterations |
|---|---|---|---|
| Fides BFGS | 41 | 6.06e+02 | 665.3 |
| Fides FIM | 146 | 1.26e+02 | 164.8 |
| PyPesto Fides | 65 | 4.51e+02 | 406.0 |
| PyPesto Fides BFGS | 51 | 7.61e+02 | 673.5 |
| PyPesto Fides FIM | 141 | 1.51e+02 | 165.4 |

**Figure 8: Parameter estimation results for the Bachmann model. (a)** Run times for each optimization setting sorted based on run time. The line denotes the median. **(b)** Waterfall plot for each optimizer. The y-axis is translated such that the minimum value equals 1. **(c)** Normalized run time versus actual run time for each optimizer option. The blue dots in each panel correspond to the optimizer option in the heading. **(d)** Summary statistics for the optimizers that converged to the best-found optima.

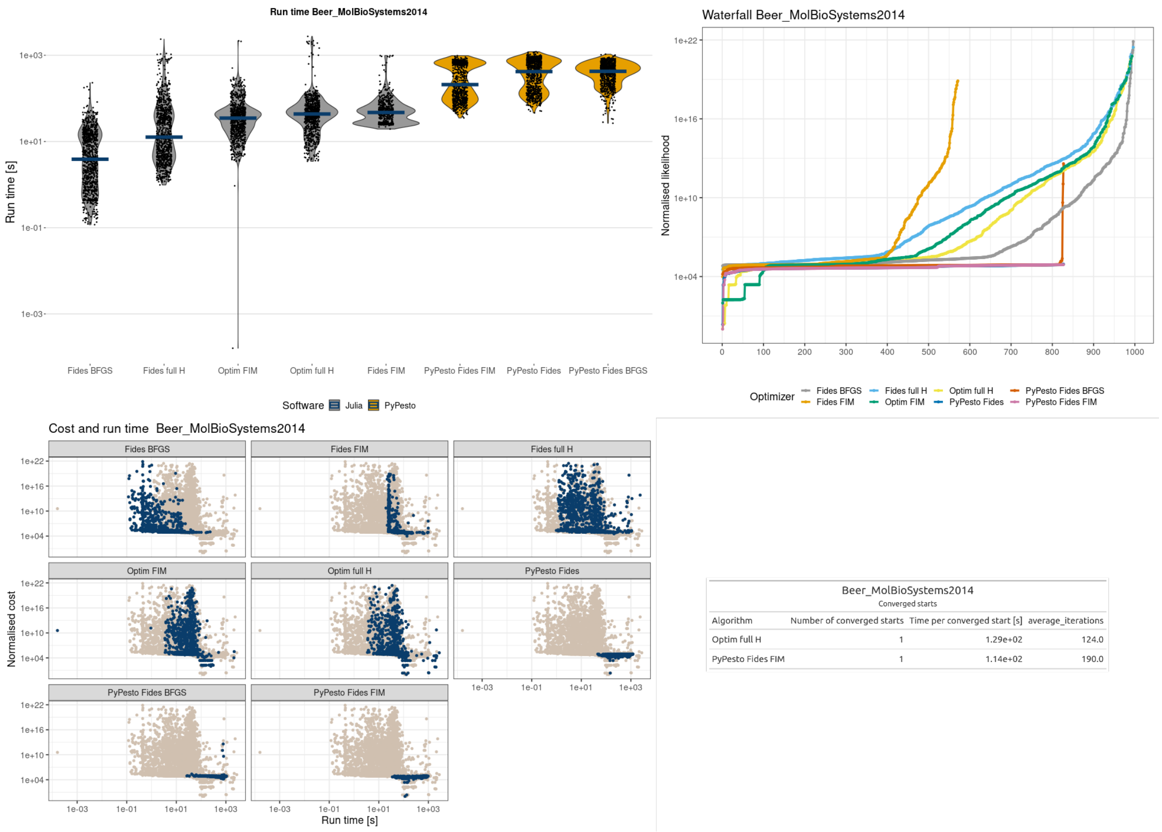


**Beer_MolBioSystems2014**

Converged starts

| Algorithm | Number of converged starts | Time per converged start [s] | average_iterations |
|---|---|---|---|
| Optim full H | 1 | 1.29e+02 | 124.0 |
| PyPesto Fides FIM | 1 | 1.14e+02 | 190.0 |

**Figure 9: Parameter estimation results for the Beer model.** **(a)**Run times for each optimization setting sorted based on run time. The line denotes the median. **(b)** Waterfall plot for each optimizer. The y-axis is translated such that the minimum value equals 1. **(c)** Normalized run time versus actual run time for each optimizer option. The blue dots in each panel correspond to the optimizer option in the heading. **(d)** Summary statistics for the optimizers that converged to the best-found optima.

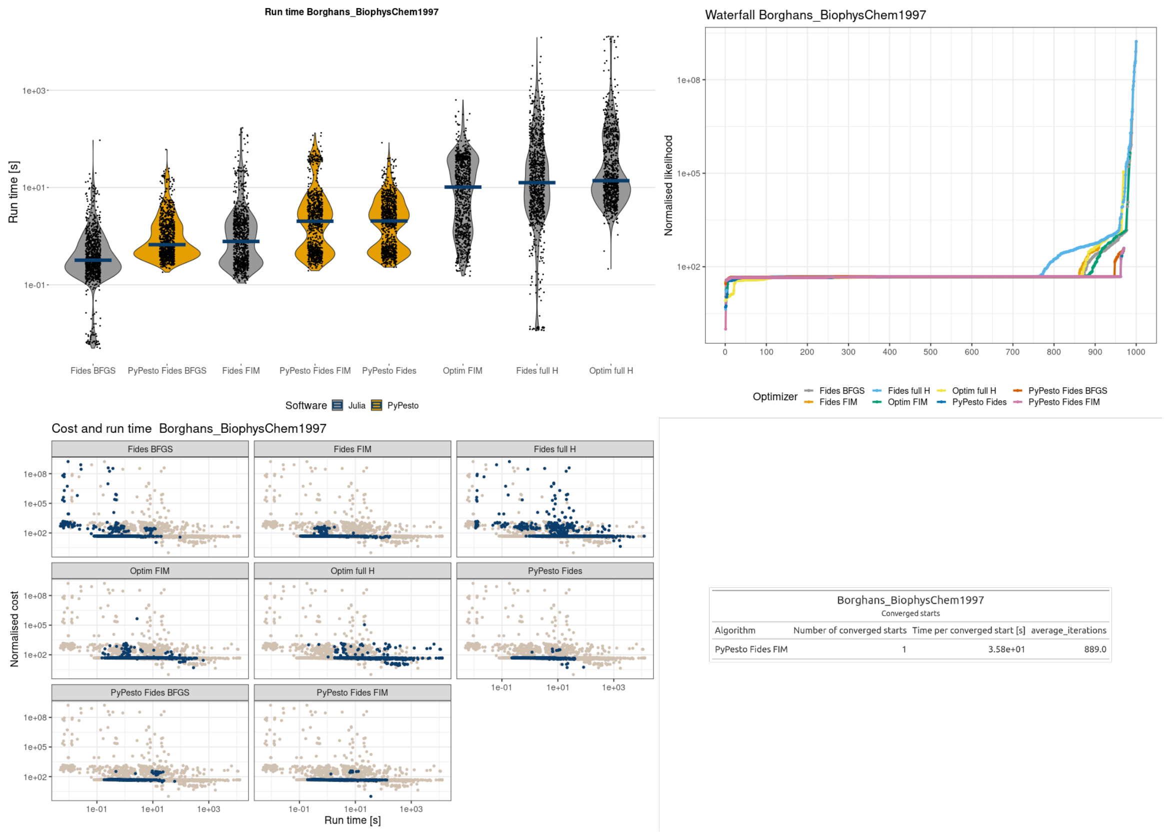


**Figure 11: Parameter estimation results for the Borghans model.** **(a)** Run times for each optimization setting are sorted based on run time. The line denotes the median. **(b)** Waterfall plot for each optimizer. The y-axis is translated such that the minimum value equals 1. **(c)** Normalized run time versus actual run time for each optimizer option. The blue dots in each panel correspond to the optimizer option in the heading. **(d)** Summary statistics for the optimizers that converged to the best-found optima.

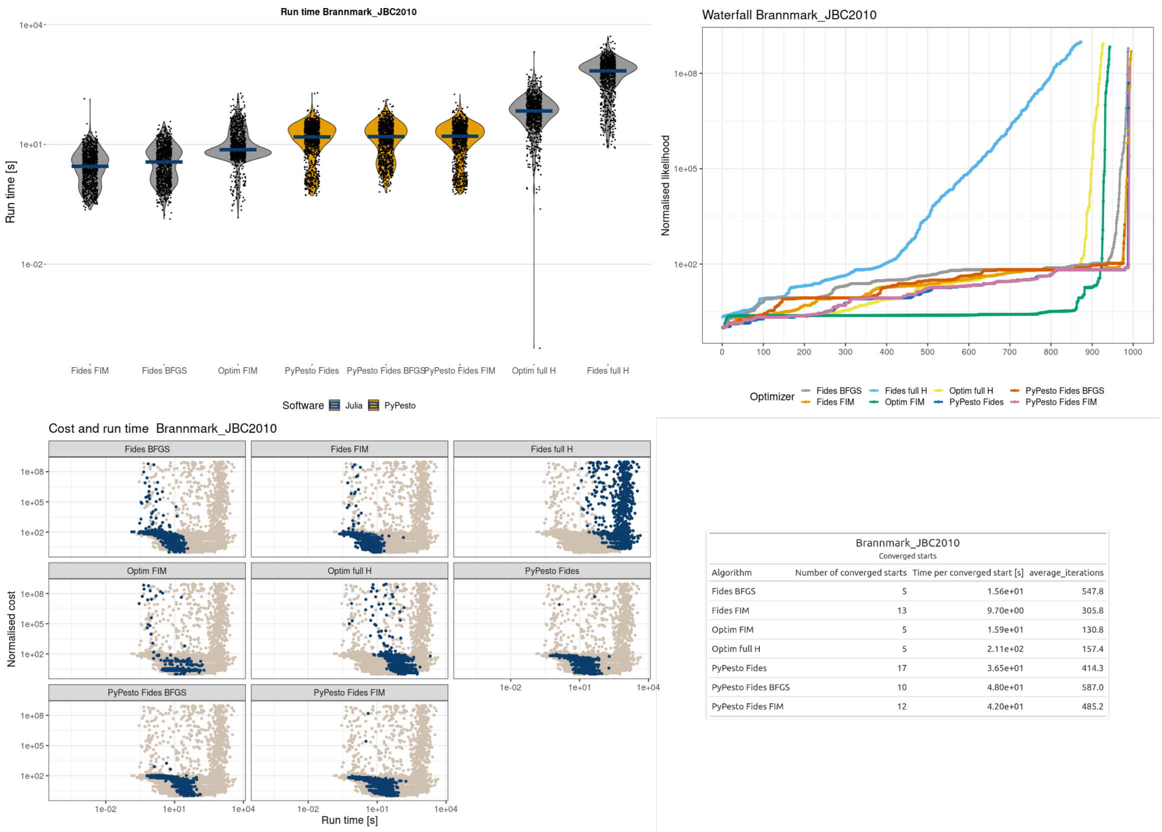


| Algorithm | Number of converged starts | Time per converged start [s] | average_iterations |
|---|---|---|---|
| Fides BFGS | 5 | 1.56e+01 | 547.8 |
| Fides FIM | 13 | 9.70e+00 | 305.8 |
| Optim FIM | 5 | 1.59e+01 | 130.8 |
| Optim full H | 5 | 2.11e+02 | 157.4 |
| PyPesto Fides | 17 | 3.65e+01 | 414.3 |
| PyPesto Fides BFGS | 10 | 4.80e+01 | 587.0 |
| PyPesto Fides FIM | 12 | 4.20e+01 | 485.2 |

**Figure 12: Parameter estimation results for the Brannmark model.** **(a)**Run times for each optimization setting are sorted based on run time. The line denotes the median. **(b)** Waterfall plot for each optimizer. The y-axis is translated such that the minimum value equals 1. **(c)** Normalized run time versus actual run time for each optimizer option. The blue dots in each panel correspond to the optimizer option in the heading. **(d)** Summary statistics for the optimizers that converged to the best-found optima.

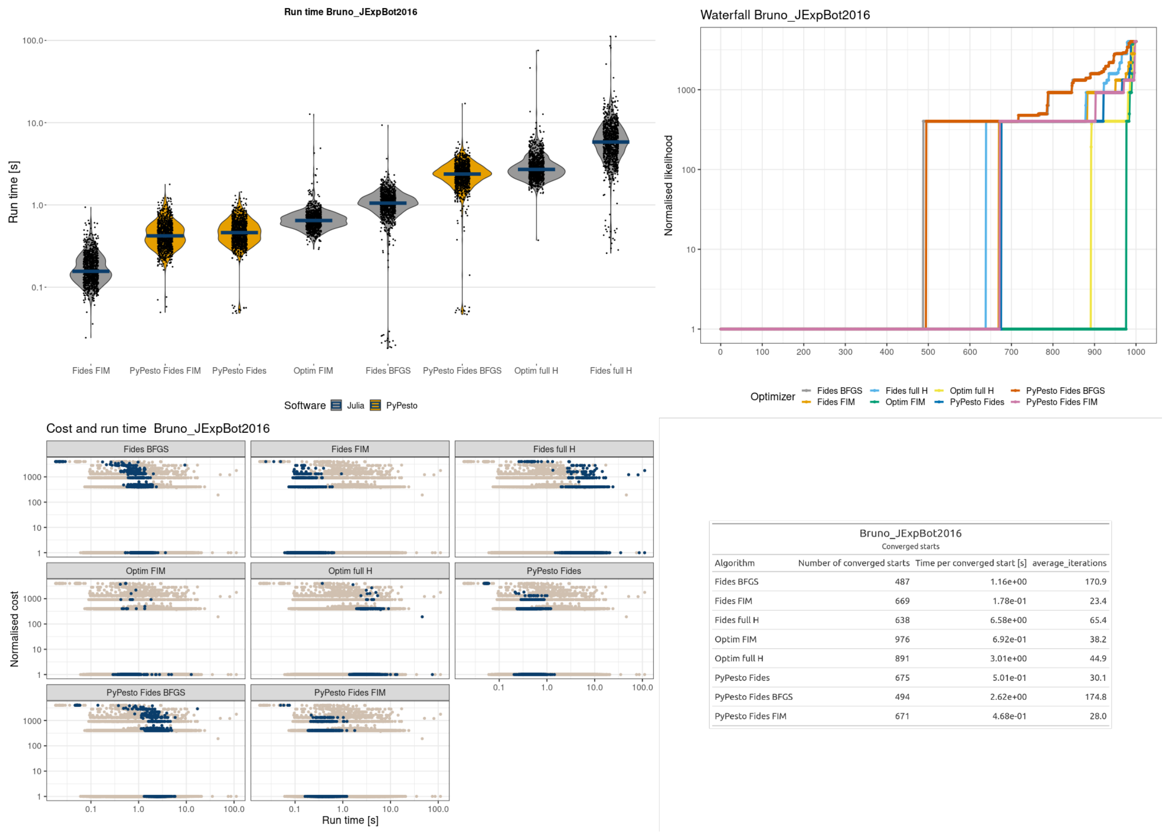


| Bruno_JExpBot2016<br>Converged starts | | | |
|---|---|---|---|
| Algorithm | Number of converged starts | Time per converged start [s] | average_iterations |
| Fides BFGS | 487 | 1.16e+00 | 170.9 |
| Fides FIM | 669 | 1.78e-01 | 23.4 |
| Fides full H | 638 | 6.58e+00 | 65.4 |
| Optim FIM | 976 | 6.92e-01 | 38.2 |
| Optim full H | 891 | 3.01e+00 | 44.9 |
| PyPesto Fides | 675 | 5.01e-01 | 30.1 |
| PyPesto Fides BFGS | 494 | 2.62e+00 | 174.8 |
| PyPesto Fides FIM | 671 | 4.68e-01 | 28.0 |

**Figure 13: Parameter estimation results for the Bruno model.** **(a)**Run times for each optimization setting are sorted based on run time. The line denotes the median. **(b)** Waterfall plot for each optimizer. The y-axis is translated such that the minimum value equals 1. **(c)** Normalized run time versus actual run time for each optimizer option. The blue dots in each panel correspond to the optimizer option in the heading. **(d)** Summary statistics for the optimizers that converged to the best-found optima.

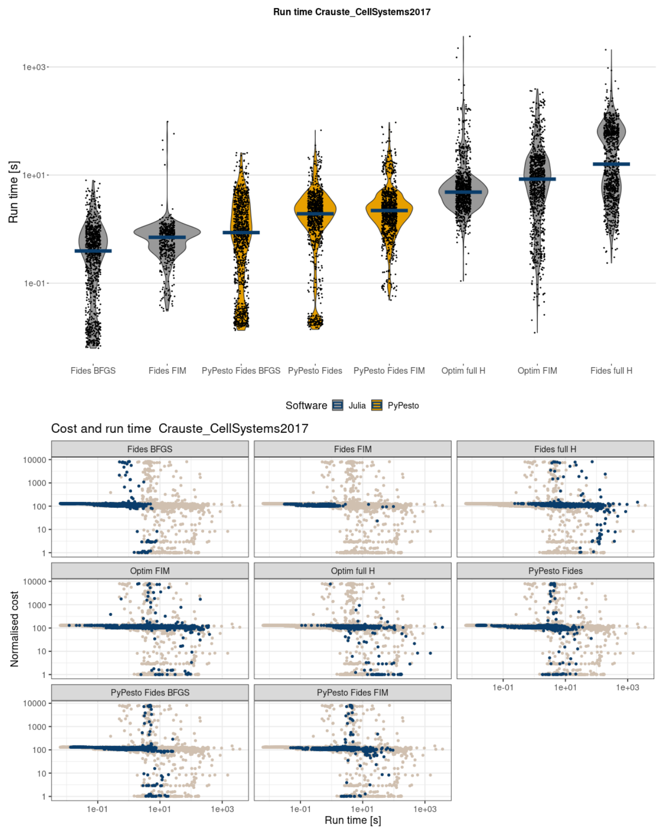


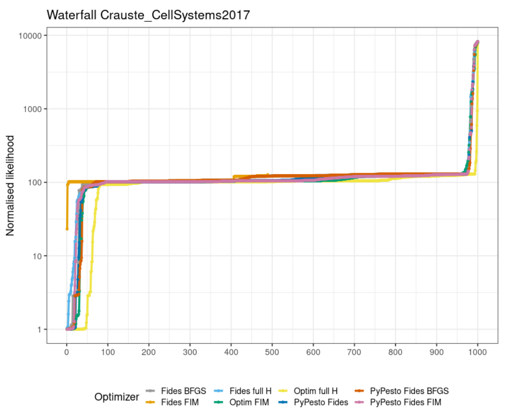


Crauste_CellSystems2017

Converged starts

| Algorithm | Number of converged starts | Time per converged start [s] | average_iterations |
|---|---|---|---|
| Fides BFGS | 10 | 2.86e+00 | 403.7 |
| Fides full H | 3 | 4.69e+01 | 457.0 |
| Optim FIM | 21 | 4.30e+01 | 640.6 |
| Optim full H | 47 | 6.82e+01 | 441.8 |
| PyPesto Fides | 21 | 6.17e+00 | 328.8 |
| PyPesto Fides BFGS | 16 | 7.57e+00 | 469.1 |
| PyPesto Fides FIM | 17 | 3.82e+00 | 234.4 |

**Figure 14: Parameter estimation results for the Crauste model.** **(a)**Run times for each optimization setting are sorted based on run time. The line denotes the median. **(b)** Waterfall plot for each optimizer. The y-axis is translated such that the minimum value equals 1. **(c)** Normalized run time versus actual run time for each optimizer option. The blue dots in each panel correspond to the optimizer option in the heading. **(d)** Summary statistics for the optimizers that converged to the best-found optima.

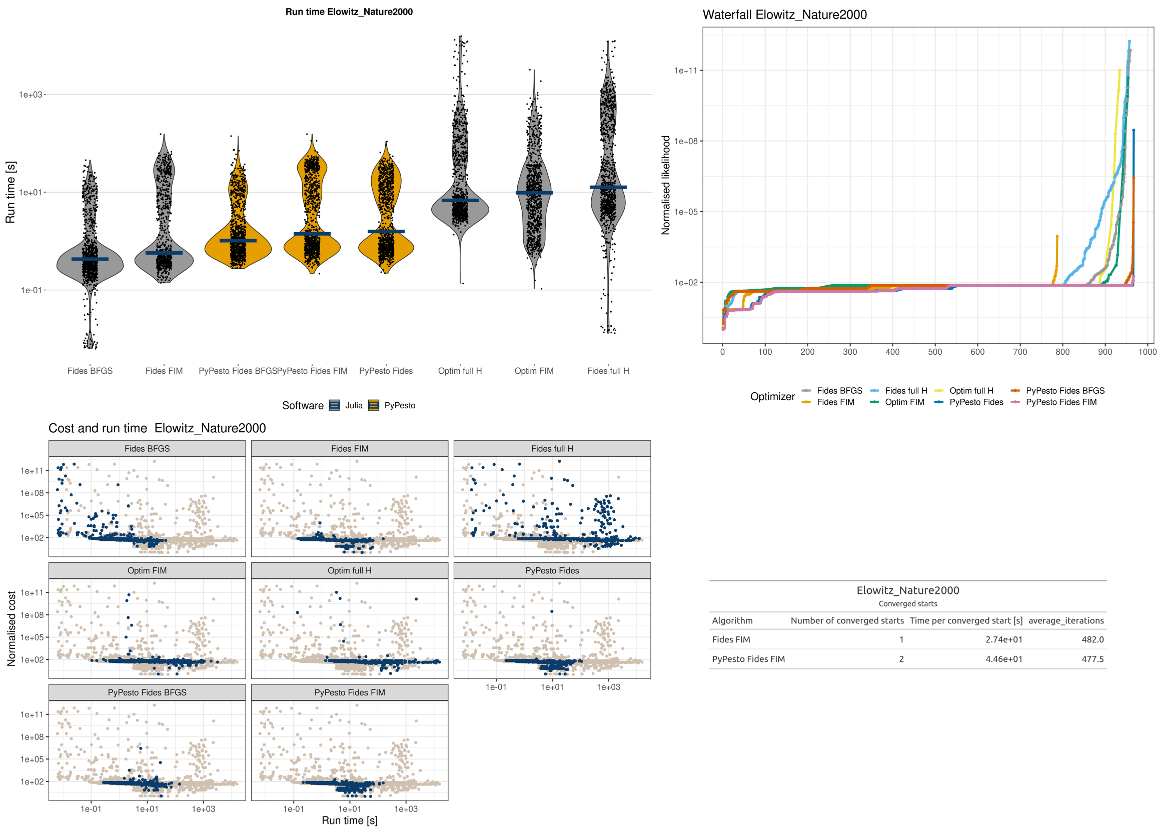


Elowitz_Nature2000
Converged starts

| Algorithm | Number of converged starts | Time per converged start [s] | average_iterations |
|---|---|---|---|
| Fides FIM | 1 | 2.74e+01 | 482.0 |
| PyPesto Fides FIM | 2 | 4.46e+01 | 477.5 |

**Figure 15: Parameter estimation results for the Elowitz model.** **(a)**Run times for each optimization setting are sorted based on run time. The line denotes the median. **(b)** Waterfall plot for each optimizer. The y-axis is translated such that the minimum value equals 1. **(c)** Normalized run time versus actual run time for each optimizer option. The blue dots in each panel correspond to the optimizer option in the heading. **(d)** Summary statistics for the optimizers that converged to the best-found optima.

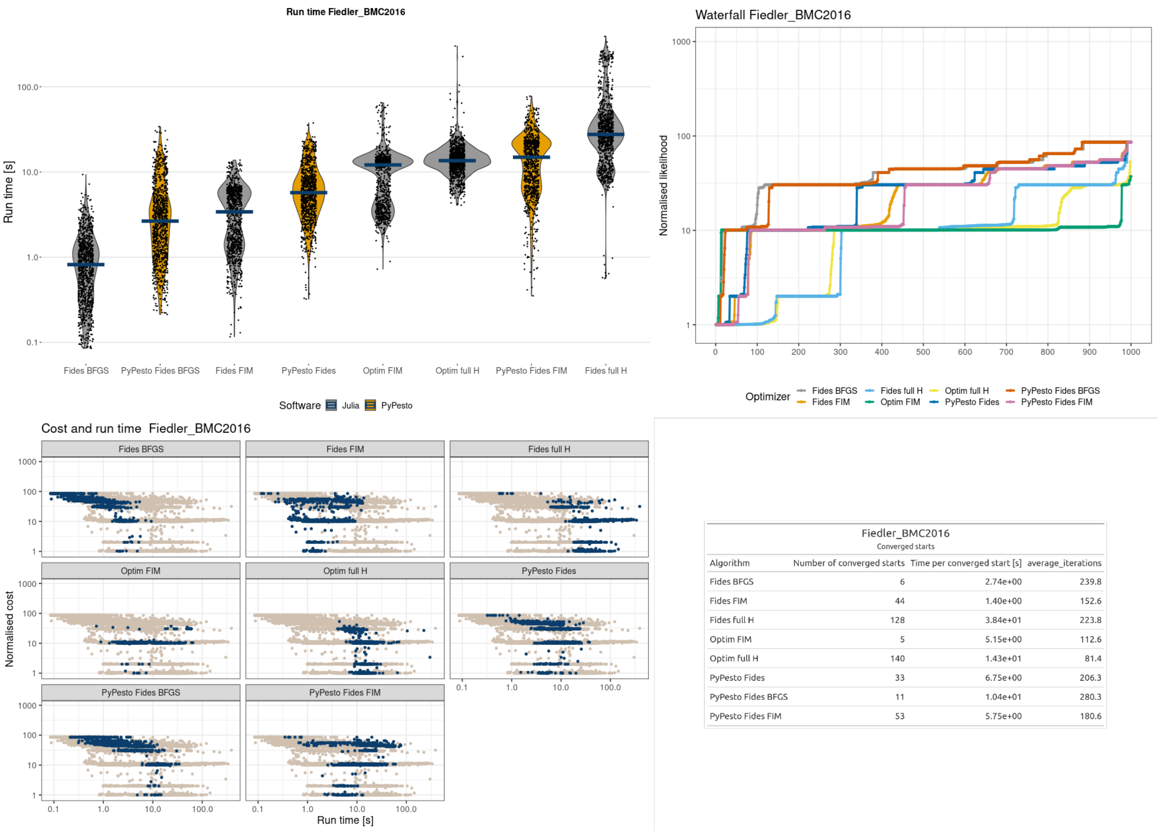


Fiedler_BMC2016

Converged starts

| Algorithm | Number of converged starts | Time per converged start [s] | average_iterations |
|---|---|---|---|
| Fides BFGS | 6 | 2.74e+00 | 239.8 |
| Fides FIM | 44 | 1.40e+00 | 152.6 |
| Fides full H | 128 | 3.84e+01 | 223.8 |
| Optim FIM | 5 | 5.15e+00 | 112.6 |
| Optim full H | 140 | 1.43e+01 | 81.4 |
| PyPesto Fides | 33 | 6.75e+00 | 206.3 |
| PyPesto Fides BFGS | 11 | 1.04e+01 | 280.3 |
| PyPesto Fides FIM | 53 | 5.75e+00 | 180.6 |

**Figure 16: Parameter estimation results for the Fiedler model.** **(a)**Run times for each optimization setting are sorted based on run time. The line denotes the median. **(b)** Waterfall plot for each optimizer. The y-axis is translated such that the minimum value equals 1. **(c)** Normalized run time versus actual run time for each optimizer option. The blue dots in each panel correspond to the optimizer option in the heading. **(d)** Summary statistics for the optimizers that converged to the best-found optima.

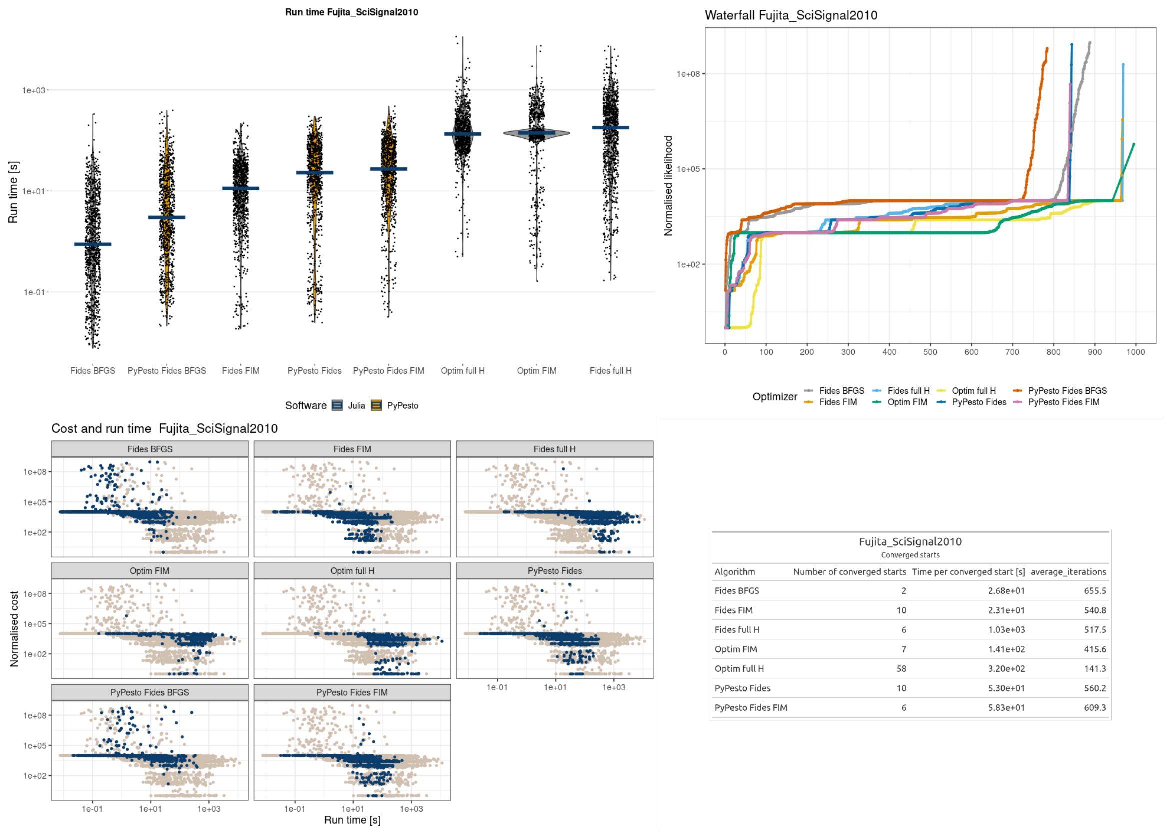


| Fujita_SciSignal2010<br>Converged starts | | | |
|---|---|---|---|
| Algorithm | Number of converged starts | Time per converged start [s] | average_iterations |
| Fides BFGS | 2 | 2.68e+01 | 655.5 |
| Fides FIM | 10 | 2.31e+01 | 540.8 |
| Fides full H | 6 | 1.03e+03 | 517.5 |
| Optim FIM | 7 | 1.41e+02 | 415.6 |
| Optim full H | 58 | 3.20e+02 | 141.3 |
| PyPesto Fides | 10 | 5.30e+01 | 560.2 |
| PyPesto Fides FIM | 6 | 5.83e+01 | 609.3 |

**Figure 17: Parameter estimation results for the Fujita model.** **(a)**Run times for each optimization setting sorted based on run time. The line denotes the median. **(b)** Waterfall plot for each optimizer. The y-axis is translated such that the minimum value equals 1. **(c)** Normalized run time versus actual run time for each optimizer option. The blue dots in each panel correspond to the optimizer option in the heading. **(d)** Summary statistics for the optimizers that converged to the best-found optima.

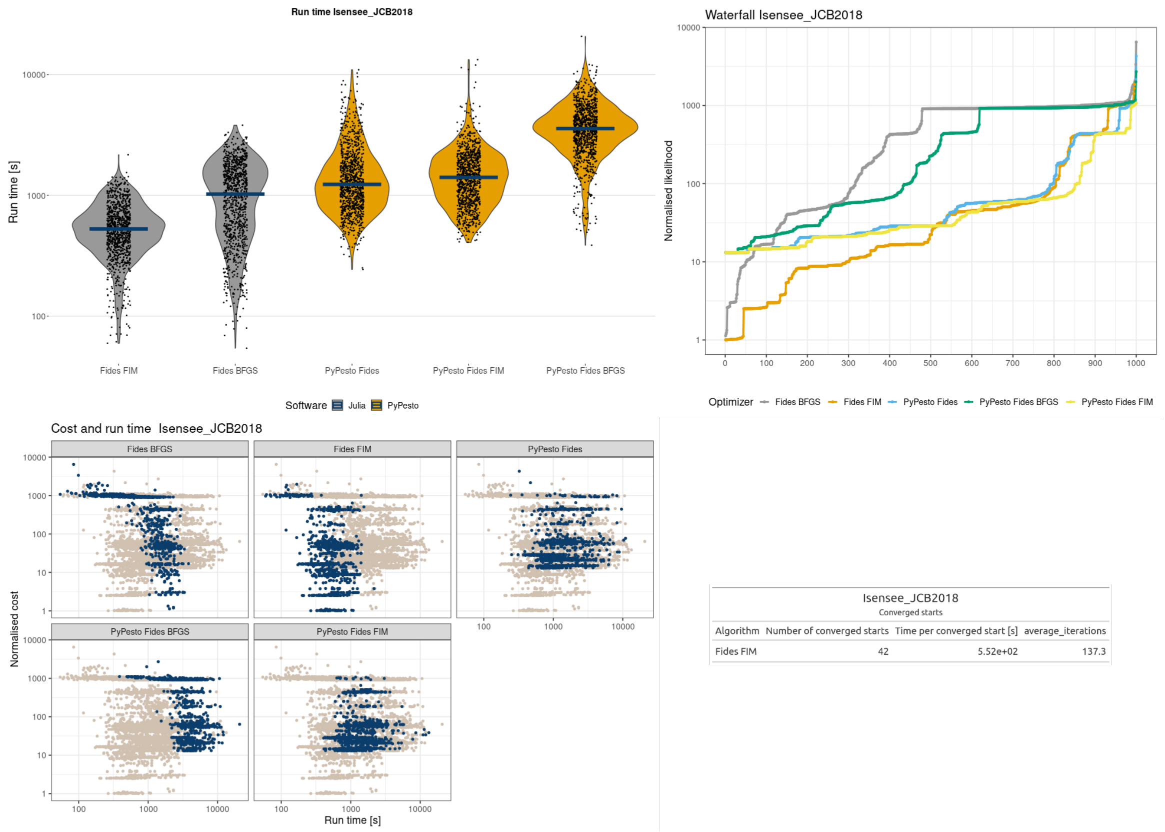


| Algorithm | Number of converged starts | Time per converged start [s] | average_iterations |
|---|---|---|---|
| Fides FIM | 42 | 5.52e+02 | 137.3 |

**Figure 18: Parameter estimation results for the Isensee model.** **(a)**Run times for each optimization setting are sorted based on run time. The line denotes the median. **(b)** Waterfall plot for each optimizer. The y-axis is translated such that the minimum value equals 1. **(c)** Normalized run time versus actual run time for each optimizer option. The blue dots in each panel correspond to the optimizer option in the heading. **(d)** Summary statistics for the optimizers that converged to the best-found optima.

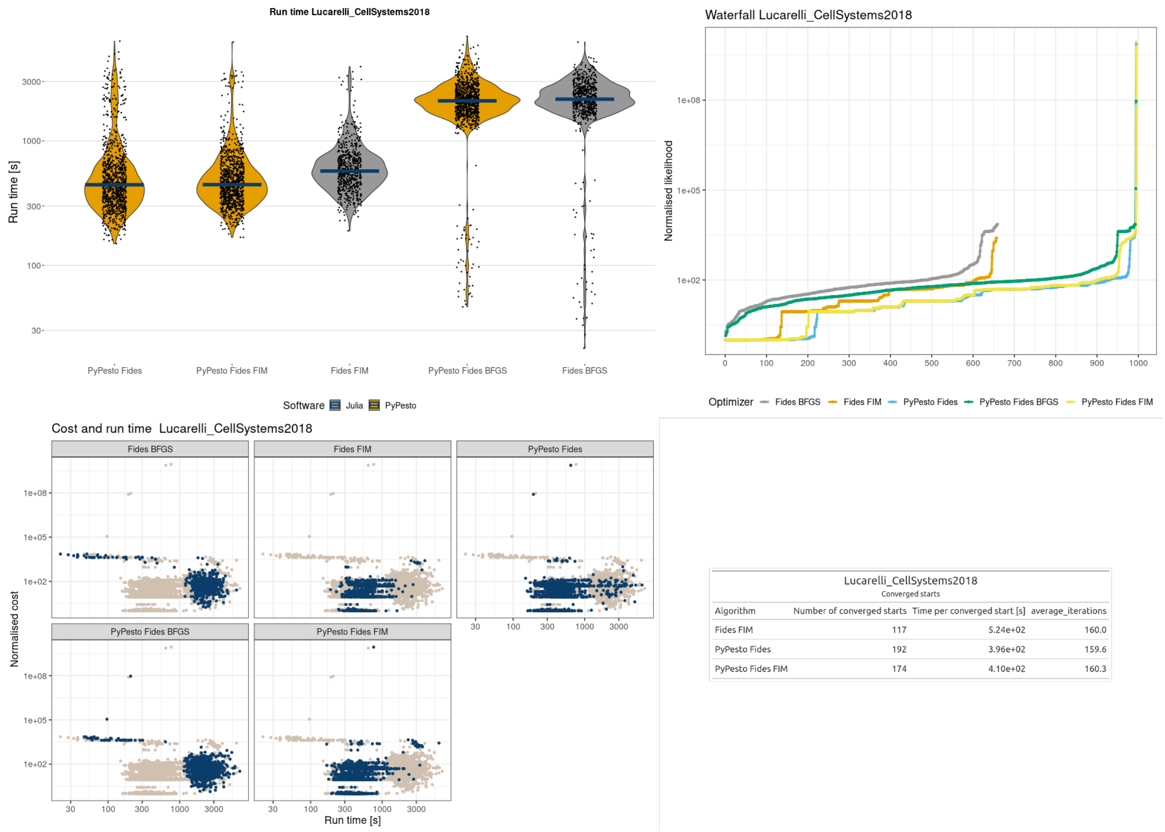


**Figure 19: Parameter estimation results for the Lucarelli model.** **(a)**Run times for each optimization setting are sorted based on run time. The line denotes the median. **(b)** Waterfall plot for each optimizer. The y-axis is translated such that the minimum value equals 1. **(c)** Normalized run time versus actual run time for each optimizer option. The blue dots in each panel correspond to the optimizer option in the heading. **(d)** Summary statistics for the optimizers that converged to the best-found optima.

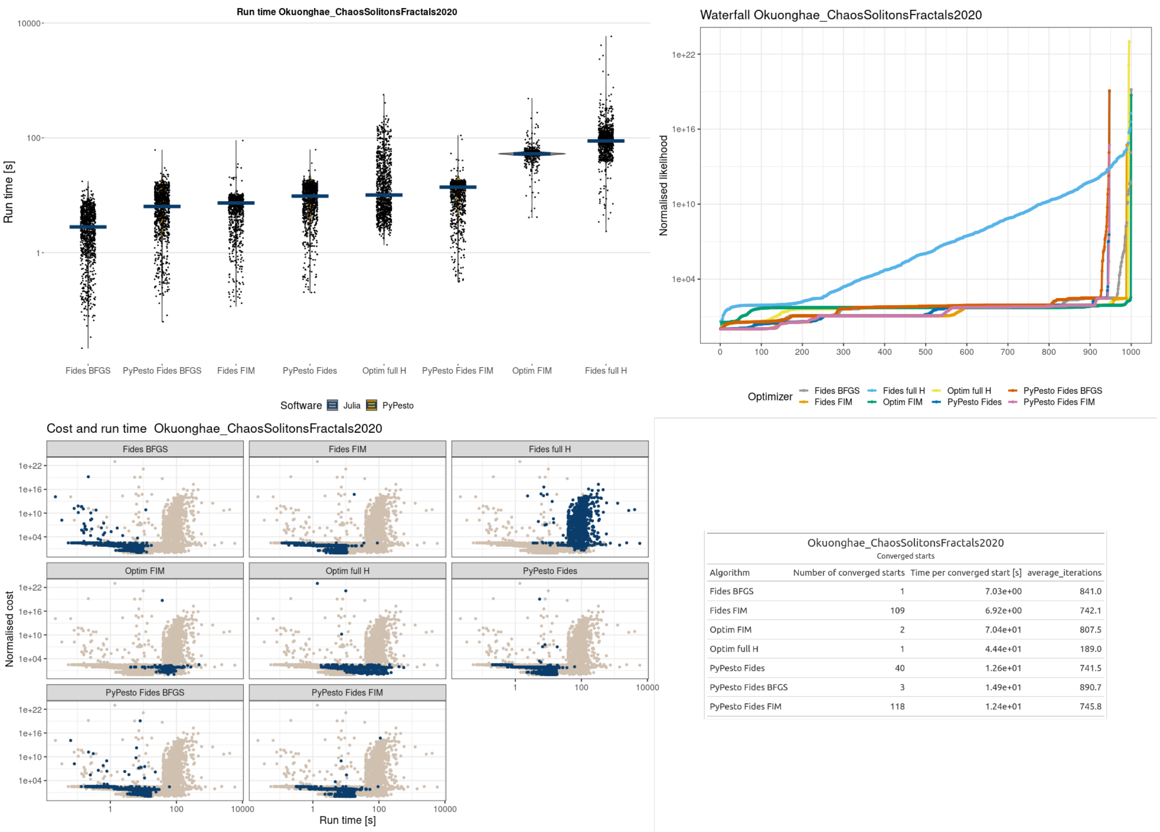


| Algorithm | Number of converged starts | Time per converged start [s] | average_iterations |
|---|---|---|---|
| Fides BFGS | 1 | 7.03e+00 | 841.0 |
| Fides FIM | 109 | 6.92e+00 | 742.1 |
| Optim FIM | 2 | 7.04e+01 | 807.5 |
| Optim full H | 1 | 4.44e+01 | 189.0 |
| PyPesto Fides | 40 | 1.26e+01 | 741.5 |
| PyPesto Fides BFGS | 3 | 1.49e+01 | 890.7 |
| PyPesto Fides FIM | 118 | 1.24e+01 | 745.8 |

**Figure 20: Parameter estimation results for the Okuonghae model.** **(a)**Run times for each optimization setting are sorted based on run time. The line denotes the median. **(b)** Waterfall plot for each optimizer. The y-axis is translated such that the minimum value equals 1. **(c)** Normalized run time versus actual run time for each optimizer option. The blue dots in each panel correspond to the optimizer option in the heading. **(d)** Summary statistics for the optimizers that converged to the best-found optima.

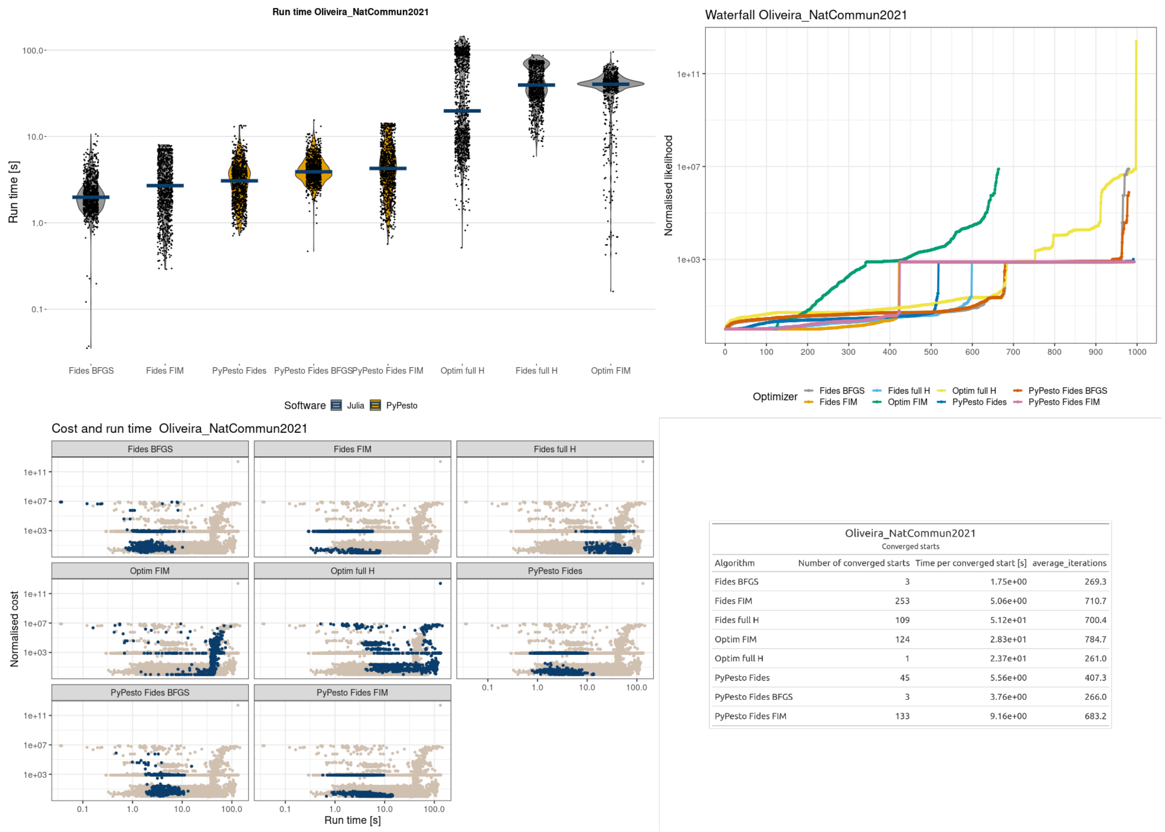


| Algorithm | Number of converged starts | Time per converged start [s] | average_iterations |
|---|---|---|---|
| Fides BFGS | 3 | 1.75e+00 | 269.3 |
| Fides FIM | 253 | 5.06e+00 | 710.7 |
| Fides full H | 109 | 5.12e+01 | 700.4 |
| Optim FIM | 124 | 2.83e+01 | 784.7 |
| Optim full H | 1 | 2.37e+01 | 261.0 |
| PyPesto Fides | 45 | 5.56e+00 | 407.3 |
| PyPesto Fides BFGS | 3 | 3.76e+00 | 266.0 |
| PyPesto Fides FIM | 133 | 9.16e+00 | 683.2 |

**Figure 21: Parameter estimation results for the Oliveira model.** **(a)**Run times for each optimization setting are sorted based on run time. The line denotes the median. **(b)** Waterfall plot for each optimizer. The y-axis is translated such that the minimum value equals 1. **(c)** Normalized run time versus actual run time for each optimizer option. The blue dots in each panel correspond to the optimizer option in the heading. **(d)** Summary statistics for the optimizers that converged to the best-found optima.

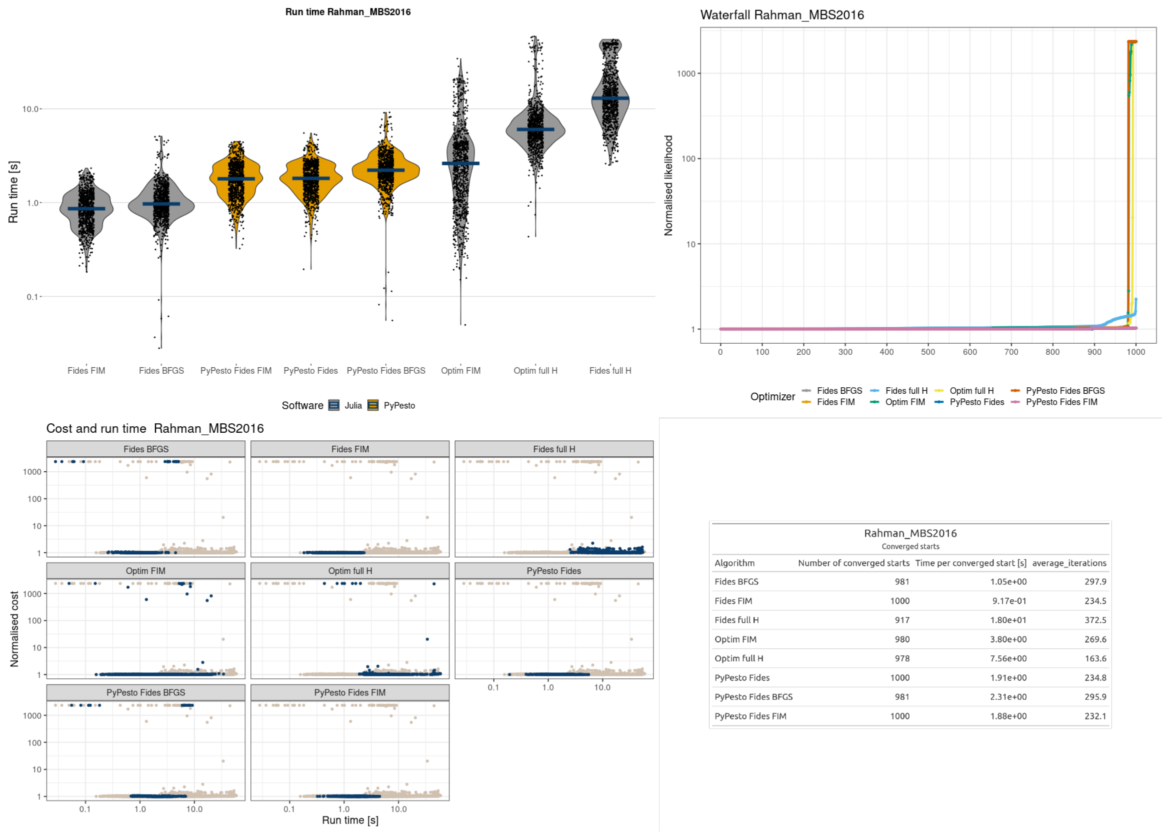


| Rahman_MBS2016<br>Converged starts | | | |
|---|---|---|---|
| Algorithm | Number of converged starts | Time per converged start [s] | average_iterations |
| Fides BFGS | 981 | 1.05e+00 | 297.9 |
| Fides FIM | 1000 | 9.17e-01 | 234.5 |
| Fides full H | 917 | 1.80e+01 | 372.5 |
| Optim FIM | 980 | 3.80e+00 | 269.6 |
| Optim full H | 978 | 7.56e+00 | 163.6 |
| PyPesto Fides | 1000 | 1.91e+00 | 234.8 |
| PyPesto Fides BFGS | 981 | 2.31e+00 | 295.9 |
| PyPesto Fides FIM | 1000 | 1.88e+00 | 232.1 |

**Figure 22: Parameter estimation results for the Rahman model.** **(a)**Run times for each optimization setting are sorted based on run time. The line denotes the median. **(b)** Waterfall plot for each optimizer. The y-axis is translated such that the minimum value equals 1. **(c)** Normalized run time versus actual run time for each optimizer option. The blue dots in each panel correspond to the optimizer option in the heading. **(d)** Summary statistics for the optimizers that converged to the best-found optima.

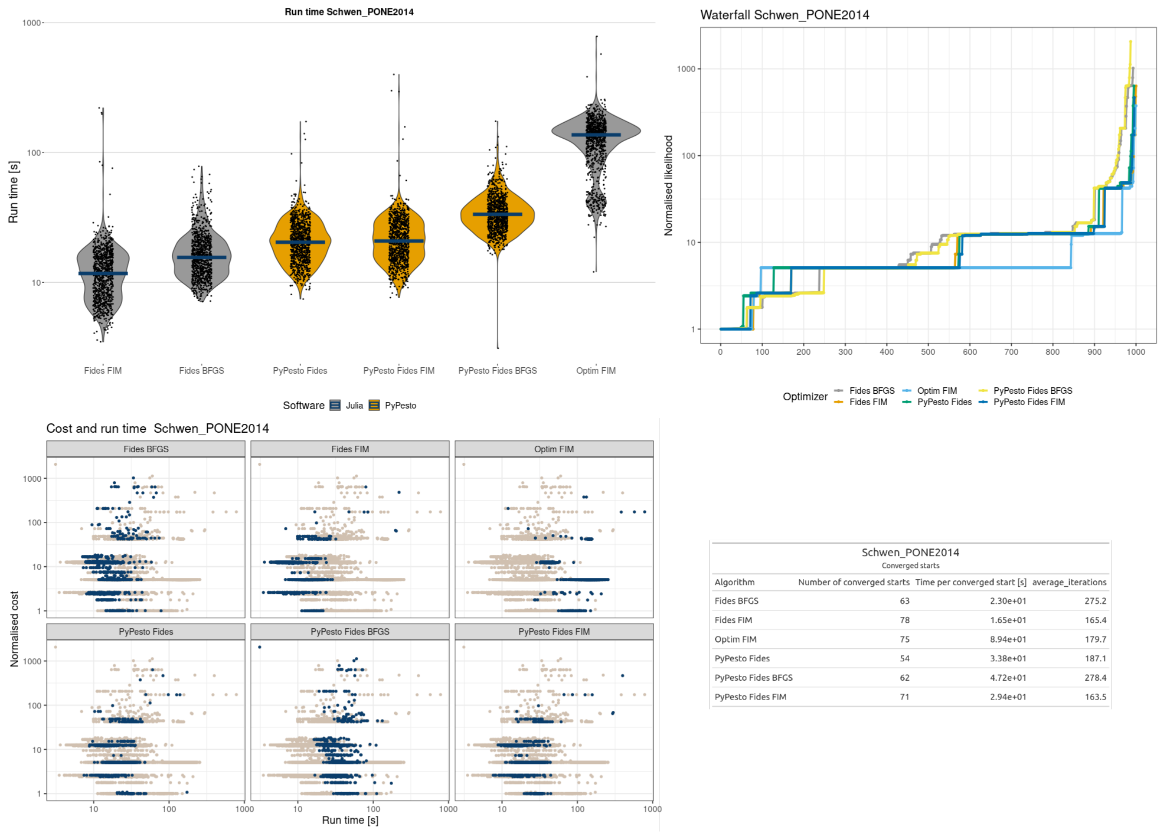


| Schwen_PONE2014<br>Converged starts | | | |
|---|---|---|---|
| Algorithm | Number of converged starts | Time per converged start [s] | average_iterations |
| Fides BFGS | 63 | 2.30e+01 | 275.2 |
| Fides FIM | 78 | 1.65e+01 | 165.4 |
| Optim FIM | 75 | 8.94e+01 | 179.7 |
| PyPesto Fides | 54 | 3.38e+01 | 187.1 |
| PyPesto Fides BFGS | 62 | 4.72e+01 | 278.4 |
| PyPesto Fides FIM | 71 | 2.94e+01 | 163.5 |

**Figure 23: Parameter estimation results for the Sneyd model.** **(a)**Run times for each optimization setting are sorted based on run time. The line denotes the median. **(b)** Waterfall plot for each optimizer. The y-axis is translated such that the minimum value equals 1. **(c)** Normalized run time versus actual run time for each optimizer option. The blue dots in each panel correspond to the optimizer option in the heading. **(d)** Summary statistics for the optimizers that converged to the best-found optima.

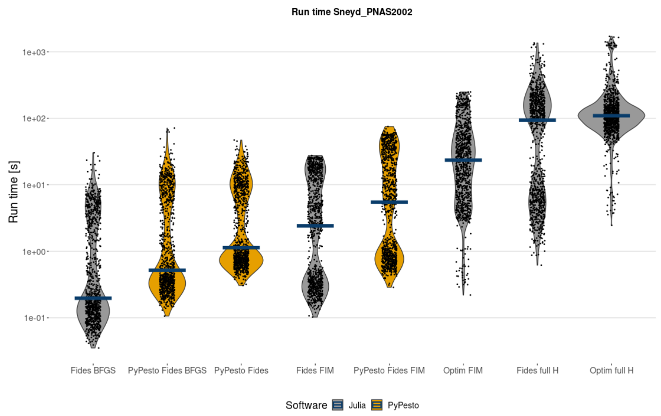


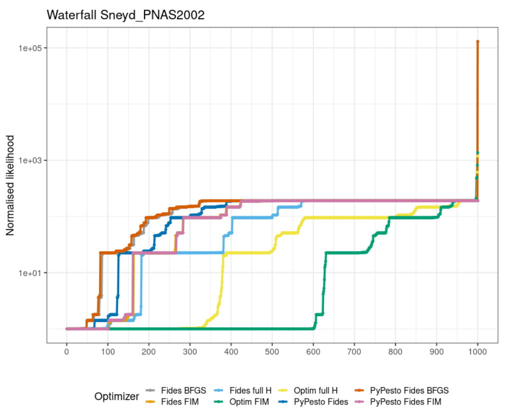


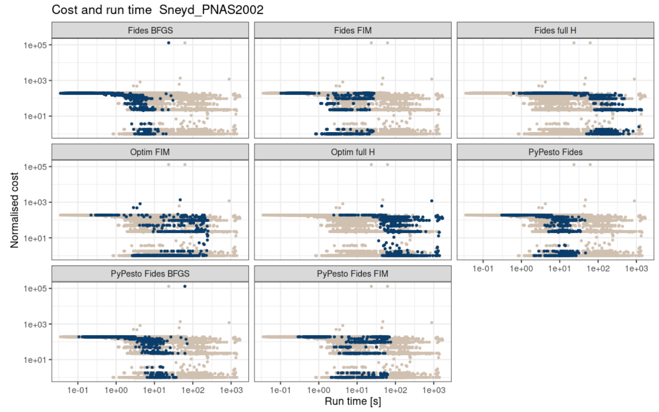


Sneyd_PNAS2002

Converged starts

| Algorithm | Number of converged starts | Time per converged start [s] | average_iterations |
|---|---|---|---|
| Fides BFGS | 48 | 6.69e+00 | 285.4 |
| Fides FIM | 103 | 5.20e+00 | 234.1 |
| Fides full H | 99 | 1.83e+02 | 154.9 |
| Optim FIM | 603 | 2.18e+01 | 171.9 |
| Optim full H | 333 | 1.81e+02 | 150.3 |
| PyPesto Fides | 67 | 9.75e+00 | 188.7 |
| PyPesto Fides BFGS | 48 | 1.48e+01 | 282.3 |
| PyPesto Fides FIM | 101 | 1.21e+01 | 239.7 |

**Figure 24: Parameter estimation results for the Sneyd model.** **(a)**Run times for each optimization setting are sorted based on run time. The line denotes the median. **(b)** Waterfall plot for each optimizer. The y-axis is translated such that the minimum value equals 1. **(c)** Normalized run time versus actual run time for each optimizer option. The blue dots in each panel correspond to the optimizer option in the heading. **(d)** Summary statistics for the optimizers that converged to the best-found optima.

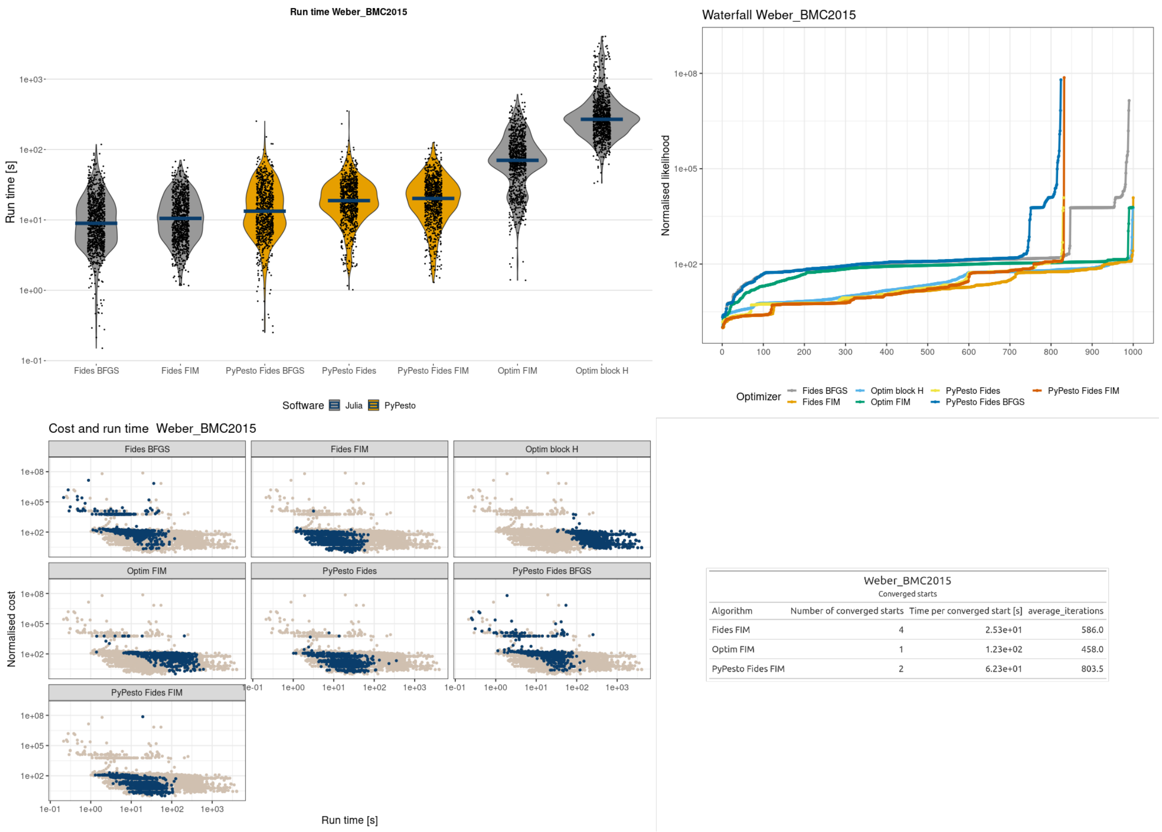


| Weber_BMC2015<br>Converged starts | | | |
|---|---|---|---|
| Algorithm | Number of converged starts | Time per converged start [s] | average_iterations |
| Fides FIM | 4 | 2.53e+01 | 586.0 |
| Optim FIM | 1 | 1.23e+02 | 458.0 |
| PyPesto Fides FIM | 2 | 6.23e+01 | 803.5 |

**Figure 25: Parameter estimation results for the Weber model.** **(a)**Run times for each optimization setting are sorted based on run time. The line denotes the median. **(b)** Waterfall plot for each optimizer. The y-axis is translated such that the minimum value equals 1. **(c)** Normalized run time versus actual run time for each optimizer option. The blue dots in each panel correspond to the optimizer option in the heading. **(d)** Summary statistics for the optimizers that converged to the best-found optima.

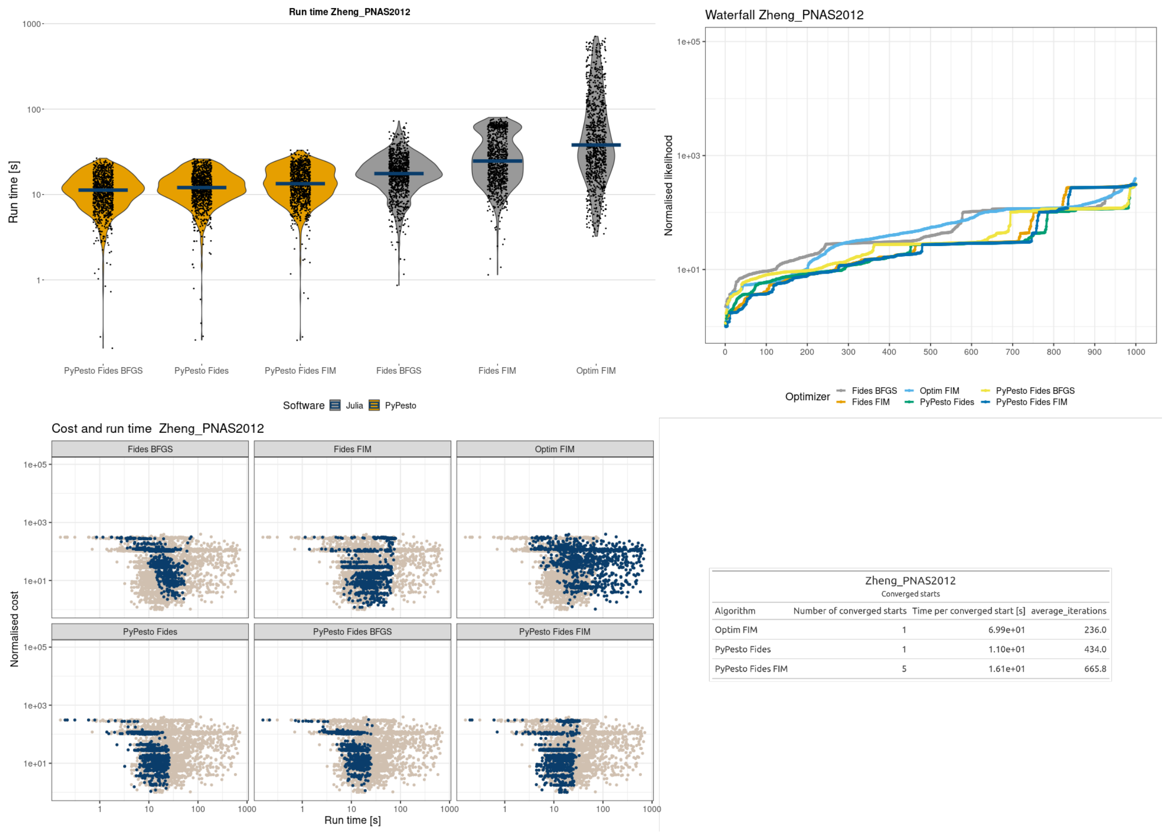


Zheng_PNAS2012

Converged starts

| Algorithm | Number of converged starts | Time per converged start [s] | average_iterations |
|---|---|---|---|
| Optim FIM | 1 | 6.99e+01 | 236.0 |
| PyPesto Fides | 1 | 1.10e+01 | 434.0 |
| PyPesto Fides FIM | 5 | 1.61e+01 | 665.8 |

**Figure 26: Parameter estimation results for the Zheng model.** **(a)**Run times for each optimization setting are sorted based on run time. The line denotes the median. **(b)** Waterfall plot for each optimizer. The y-axis is translated such that the minimum value equals 1. **(c)** Normalized run time versus actual run time for each optimizer option. The blue dots in each panel correspond to the optimizer option in the heading. **(d)** Summary statistics for the optimizers that converged to the best-found optima.
